# De novo organelle biogenesis in the cyanobacterium TDX16 released from the green alga *Haematococcus pluvialis*

**DOI:** 10.1101/161463

**Authors:** Qing-lin Dong, Xiang-ying Xing, Yang Han, Xiao-lin Wei, Shuo Zhang

**Affiliations:** Department of Bioengineering, Hebei University of Technology, Tianjin 300130, China

**Keywords:** Organelle biogenesis, Cyanobacterium, DNA acquisition, Hybridization, Transition

## Abstract

It is believed that eukaryotes arise from prokaryotes, which means that organelles can form in the latter. Such events, however, had not been observed previously. Here, we report the biogenesis of organelles in the endosymbiotic cyanobacterium TDX16 that escaped from its senescent/necrotic host cell of green alga *Haematococcus pluvialis*. In brief, organelle biogenesis in TDX16 initiated with cytoplasm compartmentalization, followed by de-compartmentalization, DNA allocation, and re-compartmentalization, as such two composite organelles-the primitive chloroplast and primitive nucleus sequestering minor and major fractions of cellular DNA respectively were formed. Thereafter, the eukaryotic cytoplasmic matrix was built up from the matrix extruded from the primitive nucleus; mitochondria were assembled in and segregated from the primitive chloroplast, whereby the primitive nucleus and primitive chloroplast matured into nucleus and chloroplast respectively; while most mitochondria turned into double-membraned vacuoles after matrix degradation. Results of pigment analyses, 16S rRNA and genome sequencing revealed that TDX16 is a phycocyanin-containing cyanobacterium resembling *Chroococcidiopsis thermalis*, which had acquired 9,017,401bp DNAs with 10301 genes form its host. Therefore, organelle biogenesis in TDX16 was achieved by hybridizing the acquired eukaryotic DNAs with its own ones and expressing the hybrid genome.

Organelle biogenesis in TDX16 results in its transition into a new eukaryotic alga TDX16-DE, which provides a reference to re-understand the development, structure, function and association of organelles in eukaryotes and the reasons behind them, and has implications on other sections of biology, particularly cancer biology and evolutionary biology: (1) the formation and maturation of the small organelle-less nascent cancer cells share striking similarities with TDX16 development and transition, so, it is most likely that cancer cells arise from bacteria; (2) organelle biogenesis in TDX16 uncovers a way of new organelle and new single-celled eukaryote formation, and in light of which, the ancestral organelles were likely formed in rather than transformed form the endosymbiotic prokaryotes that had acquired their hosts’ DNAs.

## Introduction

All cells are structurally categorized into two groups: eukaryotic cells and prokaryotic cells. Eukaryotic cells contain membrane-bounded organelles. These organelles were once thought to develop only by fission of the preexisting ones, while recent studies show that Golgi apparatus [1], peroxisomes [2–5], lysosomes [6–8] and vacuoles [9–10] form de novo. By contrast, prokaryotic cells have no organelle, but are believed to be the ancestors of eukaryotic cells, which means that organelles can develop from scratch in the former. Such events, however, had not been observed previously for unknown reasons. It is possible that organelle biogenesis in prokaryotic cells occur in specific situations and finish in short time, which result in sudden transition of the prokaryotic cells into eukaryotic cells, and thus are hard to capture.

In our previous studies we found unexpectedly that the senescent/necrotic cells of unicellular green alga *Haematococcus pluvialis* (eukaryote) suddenly burst and liberated countless small blue endosymbiotic cyanobacterial cells (TDX16) in the adverse conditions of high temperature and low irradiance (Fig. S1) [11].

Transmission electron microscopic observations revealed that tiny premature TDX16 cells with unique electron-dense heterogenous globular bodies (HGBs) multiplied by asymmetric division within the enclosing sheaths (sporangia) in the senescent/necrotic *H. pluvialis* cell (host) and subsequently grew up into small thylakoids-less endospore-producing TDX16 cells filling up the dead host’s cellular space (Fig. S2). The liberated small blue TDX16 cells are relatively stable in the dim light, but turn readily into small green algal cells as light intensity elevated [12]. The time required for TDX16’s transition is short and negatively related to light intensity, which is about 10 days at 60 μmol photons m^−2^ s^−1^. Whereas irradiance above 60 μmol photons m^−2^ s^−1^ is lethal to TDX16, causing massive cell death.

The unprecedented transition of cyanobacterium TDX16 cells into green algal cells enable us to get a real understanding of organelle biogenesis in prokaryotic cells. Hence, this research aims to study how and why organelles form in TDX16.

## Results

### Light microscopic observation of TDX16-to-alga transition

At the beginning, TDX16 cells were blue in color and reproduced by binary fission within the sporangia, containing more or less small grey vesicles (day 1, Fig. 1A). In the following day (day 2, Fig.1B) TDX16 cells turned pale blue-green and began to escape from the ruptured sporangia leaving remnant sheaths (ruptured sporangia). With time, all TDX16 cells escaped from the sporangia, which no longer divided but enlarged, rounded up and turned a little bit more blue, while the intracellular grey vesicles became nearly indistinguishable (Fig. 1C, day 3). Subsequently, TDX16 cells turned slightly yellow, in which small white vesicles began to form (Fig. 1D, day 4) and increase in number and size (Fig. 1E, day 5). As such most TDX16 cells contained several small or medium-sized white vesicles, and occasionally a newly-formed small blue vesicle (Fig. F, day 6). Radical changes occurred on day 7 (Fig. 1G): TDX16 cells turned green, in which large blue vesicles and large white vesicles formed. Thereafter, the large blue and white vesicles shrank and disappeared in TDX16 cells, while parietal chloroplasts similar to those of green algae developed in some of the last type cells (day 8, Fig. 1H). When the large blue and white vesicles disappeared completely, chloroplasts formed in most TDX16 cells but were still under development in the rest ones (day 9, Fig. 1I). Ultimately, all TDX16 cells turned into green algal cells (TDX16-DE), containing one single giant chloroplast and multiplying by formation of autospores (autosporulation) (day 10, Fig. J). Such that, the color of TDX16 cultures changed from blue (day1) to green (day 10) (Fig. 1K).

**Figure 1.**
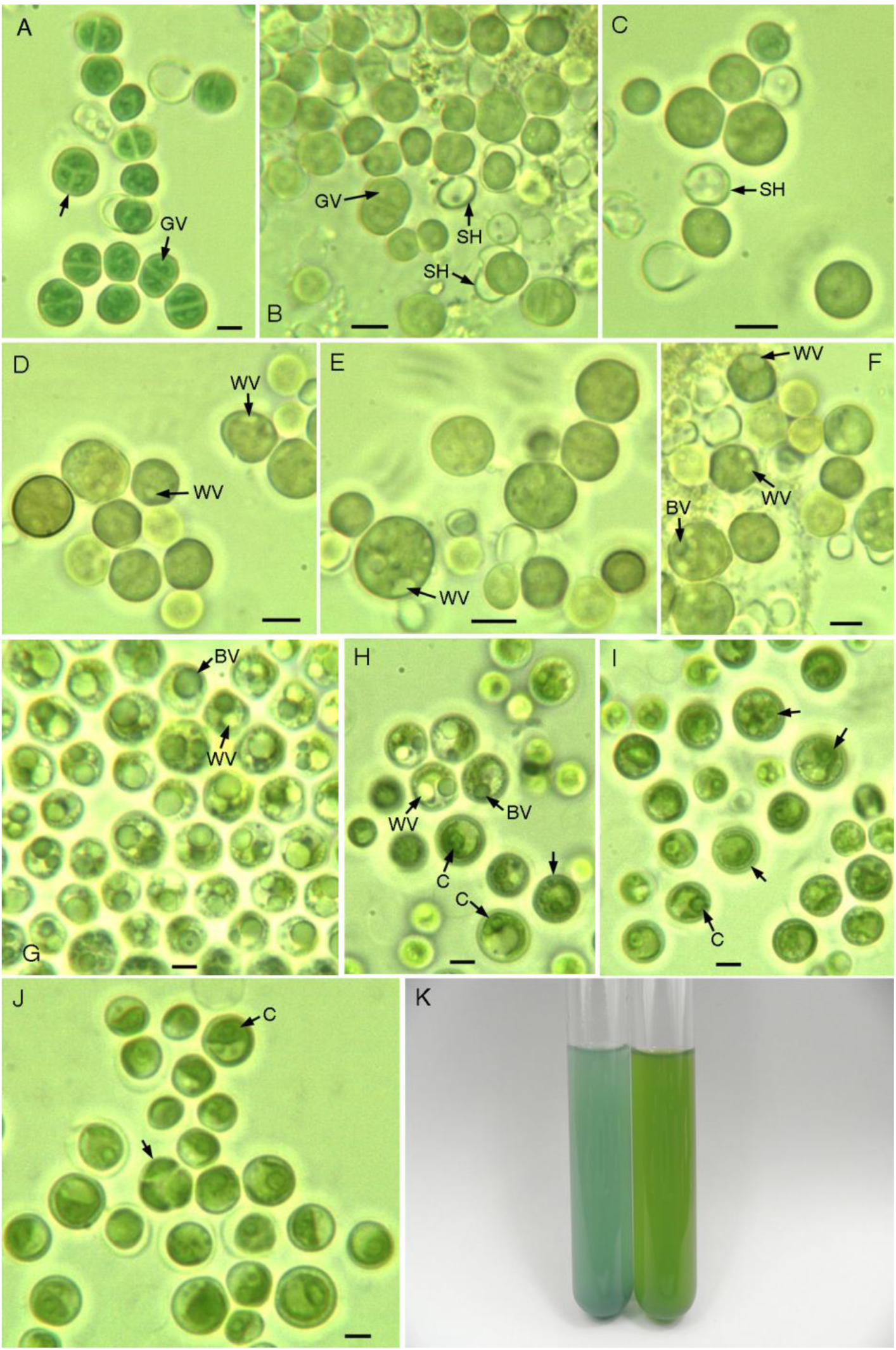
Light microscopic images of cyanobacterium TDX16-to-alga transition. **(A)** The newly inoculated blue TDX16 cells reproduced by binary fission within the sporangia (arrow) containing small grey vesicles (GV) (day 1). **(B)** TDX16 cells turned pale blue-green, some of which escaped from the sporangia leaving the remnant sheath (SH) (day 2). **(C)** All TDX16 cells escaped from the sporangia, increased in size, rounded up and turned a little bit more blue (day 3). **(D)** TDX16 cells turned slightly yellow, in which white vesicle (WV) began to form (day 4), and increased in size and number **(E)** (day 5). **(F)** TDX16 cells contained several small or one medium-sized white vesicles and occasionally a newly-formed small blue vesicles (BV) (day 6). **(G)** TDX16 cells turned green, in which large blue vesicles and large white vesicles formed (day7). **(H)** Large blue vesicles and large white vesicles shrunk in part of TDX16 cells and vanished in the rest ones (arrow), and in some of the latter type cells parietal chloroplasts (C) developed (day 8). **(I)** Chloroplasts formed in most TDX16 cells, but were under development in other ones (arrow) (day 9). **(J)** All TDX16 cells turned into green algal cells (TDX16-DE), containing single chloroplast and reproducing by autosporulation (arrow) in day 10. **(K)** Image of TDX16 culture samples on day 1(left) and day 10 (right). Scale bar 2μm.

### Transmission electron microscopic observation of organelle biogenesis

Consistent with our previous studies, light microscopic observation (Fig.1) showed unequivocally the transition of TDX16 cells into green algal cells as evidenced by the changes of cell color, morphology and reproduction mode as well as the formation of chloroplast. There is no doubt that during the10-day transition process other organelles aside from chloroplast and compartments also formed through previously unknown intermediates and subtle changes in ultrastructure, which can not be tracked in real time, but be detected discontinuously with the conventional transmission electron microscopy (TEM). Whereas, it is a formidable task to elucidate the complex and dynamic process of organelle biogenesis based on the static TEM images. Since TDX16 cells originally contained only heterogenous globular bodies (Fig. S2), all organelles developed from scratch. That is to say, changes of TDX16 initiated from a definite start-point and progressed irreversibly in one direction towards organelle biogenesis. In this unique situation, we sequenced TEM images according to sampling time, and reconstructed the general process of organelle biogenesis based on the connection and coherence of intermediates and changes in ultrastructure. However, the time for each change could not be determined owing to the unsynchronized states of cells, and thus time intervals were showed in the figure legends.

### Initial structure of TDX16 cells

Initially, TDX16 cells (day 1) were surrounded by thick sheaths and enclosed within the sporangia (Fig.2-3), resembling the endospores of thermophilic cyanobacterium *Chroococcidiopsis* sp. [13–14]. TDX16 in the same or different sporangia remained at different states with some different inclusions (Fig.2-3). As shown in Fig.2, three different-sized cells in a multilayered sporangium contained no thylakoid but unique membrane-bounded heterogenous globular bodies (HGB) and cyanobacterial inclusions, including carboxysomes (CX) [15], polyphosphate bodies (PB) [16] and osmiophilic granules (OG) [17]. The heterogenous globular bodies sequestered DNA-like electron-dense granules and filaments (resembling compacted chromatin fibers) and situated in the nucleoids (NU), where DNA fibers (DF) [18] and ribosomes (RB) [19] scattered. The prominent difference among these cells was that some small swirly and rod-shaped electron-transparent vesicles (EV) were being developed in the left large cell. Similarly, in a five-cell-containing sporangium (Fig.3), the bottom cell contained osmiophilic granules and a polyphosphate body, while many large different-shaped electron-transparent vesicles were being developed in the three middle cells, and several thylakoid-like structures were built up in the upper cell. These results confirmed the light microscopic observations that TDX16 was hypersensitive to light and highly variable in nature, whose change could not be suppressed completely even in the dim light.

**Figure 2.**
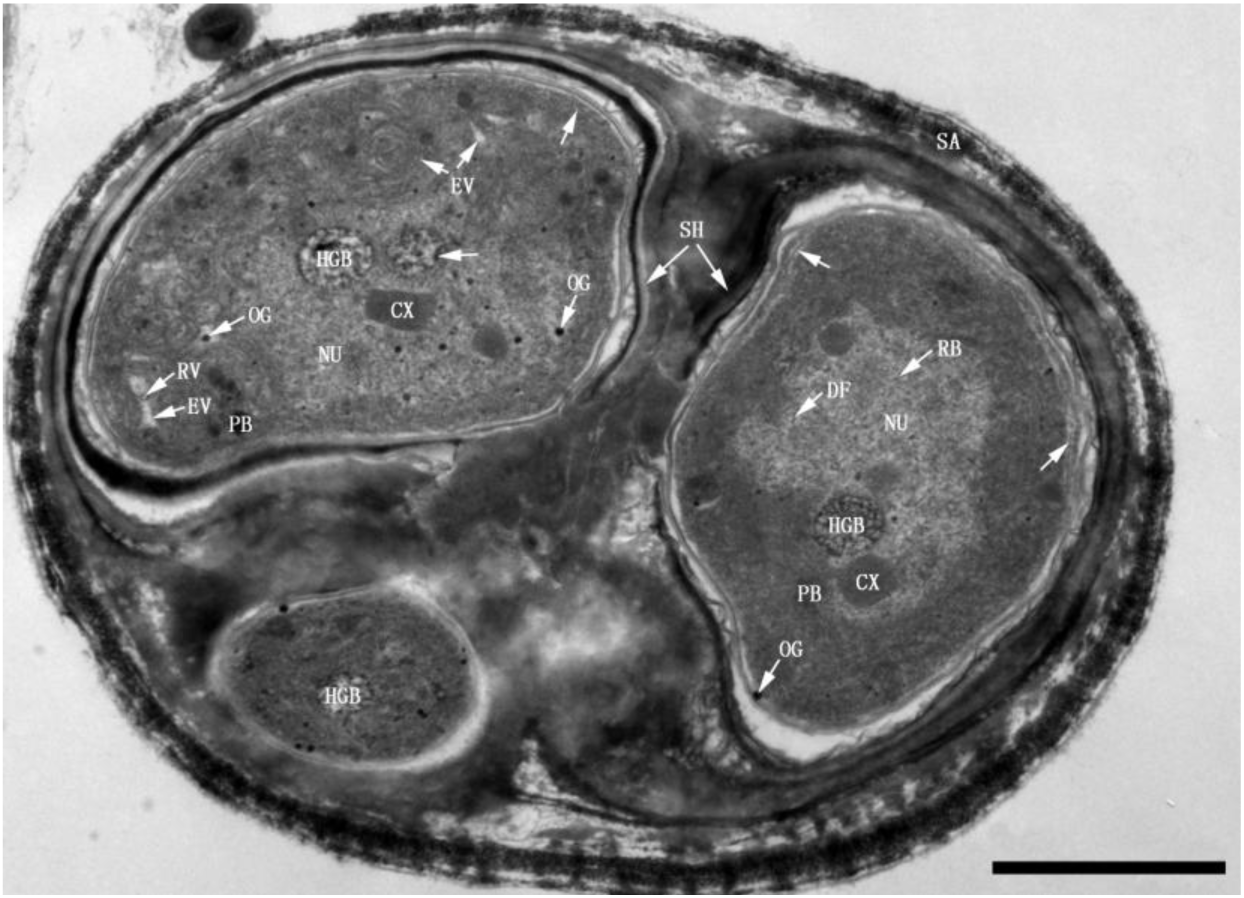
Three TDX16 cells within a sporangium (SA) (day 1). TDX16 cells were enclosed by thick sheaths (SH), containing no thylakoid, but heterogenous globular bodies (HGB), carboxysomes (CX), ribosomes (RB), DNA fibers (DF) and osmiophilic granules (OG) in the nucleoids (NU) as well as polyphosphate bodies (PB) in the cytoplasm. OG also presented in the cytoplasm and some small electron-transparent vesicles (EV) with internal ring-shaped vesicles (RV) or OG were being developed in the upper left cell. Compartmentalization initiated in the two large cells (arrow). Scale bar, 1μm.

**Figure 3.**
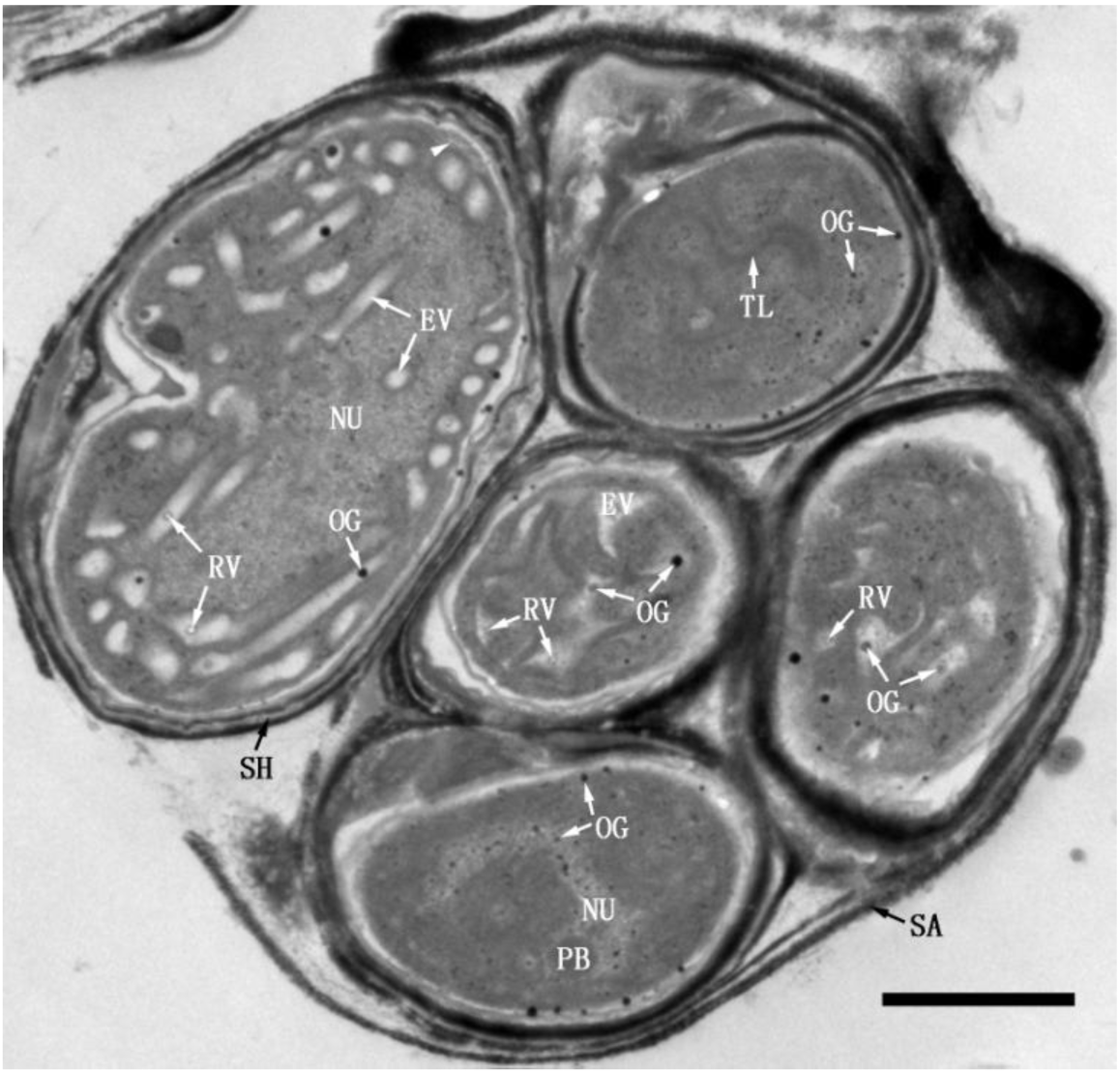
Five TDX16 cells within a sporangium (SA) (day 1). The bottom cell contained osmiophilic granules (OG), while electron-transparent vesicles (EV) were developed in the three middle cells, and several thylakoid-like structures (TL) were built up in the upper cell. Compartmentalization commenced in the large cell (arrowhead). Scale bar, 1μm.

The electron-transparent vesicles developed from osmiophilic granules, because (1) osmiophilic granule was the only membranous cyanobacterial inclusion prior to electron-transparent vesicles, (2) osmiophilic granules presented in electron-transparent vesicles, some of which were not in the section plane and thus invisible, and (3) as electron-transparent vesicles enlarged, osmiophilic granules turned into ring-shaped vesicles (RV) after their dense matrixes became opaque and finally transparent (Fig.2-3). Osmiophilic granule contains triacylglycerol and tocopherol [20], while its detail composition and structure are unknown. There is a general consensus that osmiophilic granule in cyanobacteria is comparable to plastoglobule (PG) in algal and plant chloroplasts [17, 21], which contains lipids, carotenoids, enzymes, and proteins e.g., vesicle-inducing protein in plastids 1, and structurally consists of a monolayer lipid membrane (half unit membrane) and a neutral lipid core [22–26]. Whereas, the formation of ring-shaped vesicles in the electron-transparent vesicles indicated that osmiophilic granules had two monolayer lipid membranes: the intermembrane space was likely filled with hydrophobic neutral lipids, while the interior monolayer membrane encased probably a hydrophilic “protein core”. So, as the intermembrane space dilated, the outer monolayer membrane bulged out into an electron-transparent vesicle; while the interior monolayer membrane and protein core remained unchanged as a monolayer-membrane-bounded osmiophilic granule, which subsequently transformed into a ring-shaped vesicle after metabolizing the protein core (Fig.2-3).

### Compartmentalization, formation of cytoplasmic envelope and primary thylakoids

#### Compartmentalization, production of osmiophilic granules, dilation of electron-transparent vesicles and formation of cytoplasmic envelope

##### Compartmentalization

TDX16 with surrounded sheath escaped from the ruptured sporangium and changed rapidly in structures and inclusions. As shown in Fig.4A, the cyanobacterial polyphosphate bodies and carboxysomes vanished; the heterogenous globular bodies became nearly empty leaving only few DNA-like fibrils and electron-dense margin residues. While a startling number of osmiophilic granules and stacks of membranous elements emerged, and many small electron-transparent vesicles were being developed. The cell wall became clear, which, like those of gram-negative bacteria and other cyanobacteria [27–28], was composed of an electron-dense outer membrane (OM), an electron-transparent intermediate space (layer) and an inner electron-dense peptidoglycan layer (P), and separated from the cytoplasmic membrane (CM) by an electron-transparent extracytoplasmic (periplasmic) space (ES) (Fig.4A).

**Figure 4.**
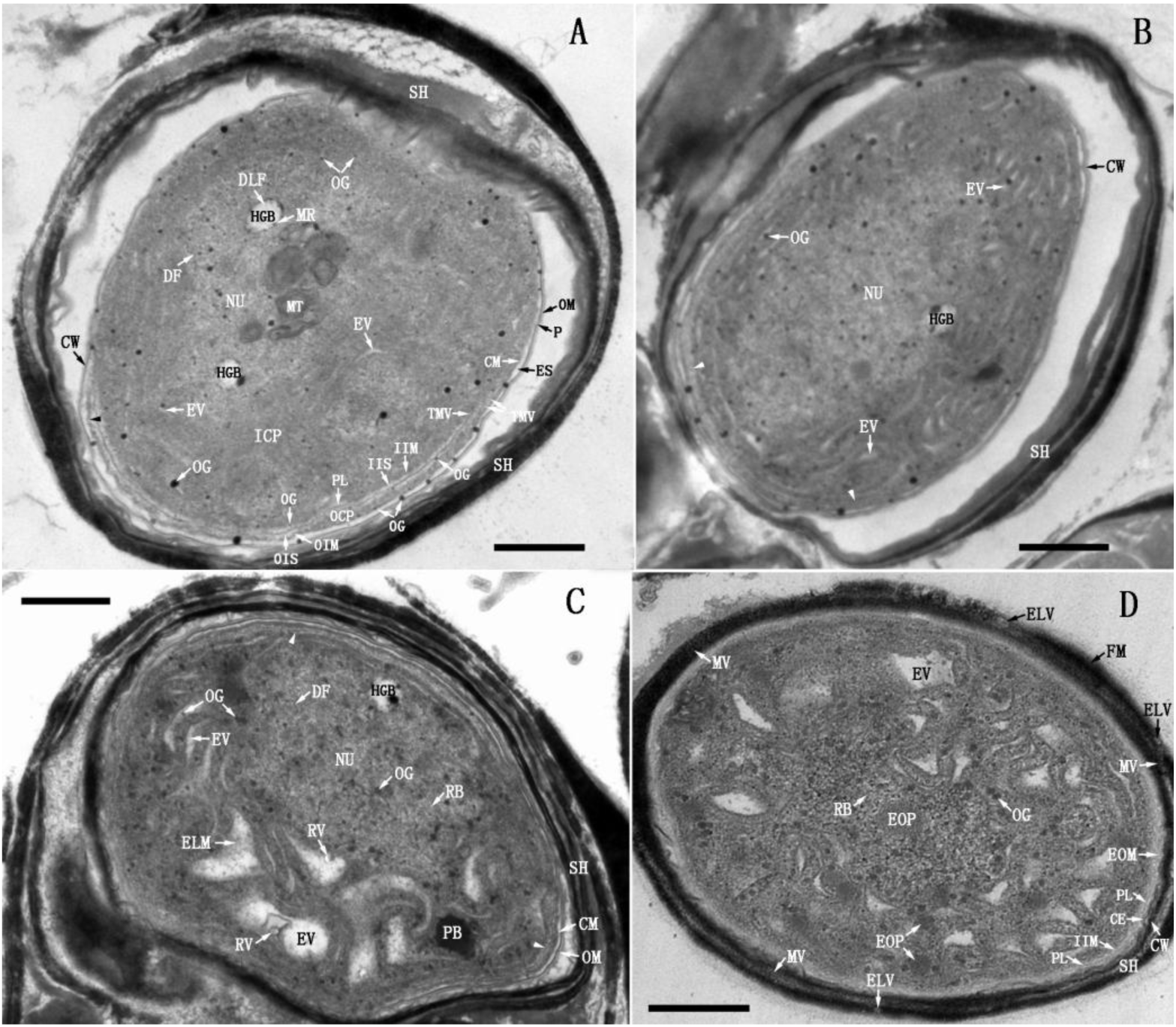
Compartmentalization, formation of electron-transparent vesicles and cytoplasmic envelope (day 1-2). (**A**) TDX16’s cell wall (CW) comprised an outer membrane (OM) and a peptidoglycan layer (P), which was separated from the cytoplasmic membrane (CM) by an extracytoplasmic space (ES). Inside the cytoplasm, an inner intracytoplasmic membrane (IIM), an outer intracytoplasmic membrane (OIM) and an intervening peptidoglycan-like layer (PL) were being synthesized by fusion of the small thick margin vesicles (TMV) blistered form CM. Whereby, the cytoplasm was partitioned into three compartments: the inner cytoplasm (ICP); the outer cytoplasm (OCP), and the sandwiched intracytoplasmic space (IS) that was further separated by PL into an outer intracytoplasmic space (OIS) and an inner intracytoplasmic space (IIS). OCP began to reduce in localized region near the start point (arrowhead), such that OIM moved to CM. Osmiophilic granules (OG) budded from CM, IIM and OIM, and migrated into ICP, where many small electron-transparent vesicles (EV) were being formed and stacks of membranous elements (MT) emerged; while the heterogenous globular bodies (HGB) became nearly empty leaving only DNA-like fibrils (DLF) and electron-dense margin residues (MR). Interestingly, OG shed from the outer leaflet of CM into ES, connecting CM and CW. (**B**) IS became narrow (arrow), while more and more small EV were being developed (**C**) EV dilated into swirling ones spiraling around the nucleoids (NU). (**D**) OCP vanished, OIM and CM were positioned together giving rise to a double-membraned cytoplasmic envelope (CE); NU and HGB disappeared, several electron-opaque particles (EOP) emerged; IS became widened filling with electron-opaque materials (EOM); a new sheath (SH) was formed, consisting of flocculent fibrillary materials (FM), microvesicles (MV) and electron-translucent vesicles (ELV). Scale bar, 0.5 μm.

The most striking change was the compartmentalization of prokaryotic cytoplasm (cytoplasm) (Fig.4A): two intracytoplasmic membranes and an intervening peptidoglycan-like layer (PL) were being synthesized synchronously, which initiated from a start point and extended parallel to the cytoplasmic membrane. As a result, the cytoplasm was partitioned into three compartments: (1) the inner cytoplasm (ICP) delimited by the inner intracytoplasmic membrane (IIM), (2) the outer cytoplasm (OCP) bounded by the outer intracytoplasmic membrane (OIM) and cytoplasmic membrane, and (3) the sandwiched intracytoplasmic space (IS) that was further separated by the peptidoglycan-like layer into an outer intracytoplasmic space (OIS) and an inner intracytoplasmic space (IIS) (Fig.4A). It was important that the outer cytoplasm began to degrade in localized regions near the start point, such that the outer intracytoplasmic membrane got closed to the cytoplasmic membrane (Fig.4A). In fact, compartmentalization also commenced in some of the newly inoculated cells (day 1, Fig.2-3). The intracytoplasmic membranes and peptidoglycan-like layer were synthesized by fusion of the small thick margin vesicles (TMV) blistered form the inner leaflet of the cytoplasmic membrane (Fig.4A) in two probable ways: (1) if the small thick margin vesicles were delimited by a half-unit membrane, they first released their contents for synthesizing the septal peptidoglycan-like layer and then fused alternately on its two sides; (2) if the small thick margin vesicles were limited by a unit membrane, as they fused one another, the septal peptidoglycan-like layer was synthesized within the coalesced vesicles, a scenario somewhat similar to the formation of cell plate during cytokinesis [29].

##### Production of osmiophilic granules

Osmiophilic granules of various sizes blistered from the inner and outer leaflets of the cytoplasmic membrane (Fig.4A), such a scenario was also observed in the newly inoculated cells (Fig.2-3). Importantly, some small osmiophilic granules also budded from the intracytoplasmic membranes (Fig. 4A). These results suggested that the intracytoplasmic membranes were functionally comparable to the cytoplasmic membrane that was apparently capable of both photosynthesis and respiration just like the cytoplasmic membrane of the thylakoid-less cyanobacterium *Gloeobacter violaceus* [30–31]. Osmiophilic granules appeared to play different roles: these blistered from the inner leaflet of cytoplasmic membrane and the intracytoplasmic membranes migrated into the inner cytoplasm for developing electron-transparent vesicles (Fig.4A); while those shed off the outer leaflet of cytoplasmic membrane contacted the cell wall, and thus served likely as transport conduits or transport vesicles to channel or transfer lipids and carotenoids from cytoplasmic membrane to cell wall (Fig.4A), because these compounds were synthesized on cytoplasmic membrane but deposited in outer membrane [32].

##### Dilation of electron-transparent vesicles and formation of cytoplasmic envelope

As the intracytoplasmic membranes and peptidoglycan-like layer extended progressively (Fig.4B) and closed up (Fig.4C), the small electron-transparent vesicles elongated (Fig.4B) and dilated asymmetrically into swirling ones spiraling around the nucleoid (Fig.4C). Thereafter, the outer cytoplasm vanished and thus the outer intracytoplasmic membrane and cytoplasmic membrane combined into a double-membraned cytoplasmic envelope (CE), which abutted the cell wall owing to the narrowing of extracytoplasmic space (Fig.4D). In parallel with these changes: (1) the heterogenous globular body and nucleoid disappeared, while electron-opaque particles (EOP) formed (Fig. 4D); (2) the intracytoplasmic space widened, filling with electron-opaque materials (EOM) (Fig. 4D); and (3) the old thick sheath scaled off, while a new fuzzy loosely compacted sheath external to the outer membrane formed, comprising of fibrillar materials (FM), microvesicles (MV) and electron-translucent vesicles (ELV).

#### Formation of primary thylakoids

As shown in Fig.5A, the intraluminal ring-shaped vesicles swelled up into dilated-ring-shaped vesicles (DRV), whose membranes ultimately met and combined with the electron-transparent vesicle membranes, giving rise to unit-membrane-bounded combined vesicles (CV); and then the combined vesicles coalesced into long ones or flattened out into short slender sacs, which were morphologically similar to cyanobacterial thylakoids, termed primary thylakoids (PT). In this way, the electron-transparent vesicles developed progressively into short primary thylakoids with opaque luminal matrix, distributing randomly in the inner cytoplasm (Fig.5B). After that, the short primary thylakoids extended or merged end-to-end into longer ones with concurrent formation of cyanophycin granules (CG) [33] and electron-opaque particles; while the long combined vesicle flattened out into primary thylakoids stepwise by localized-constriction (Fig. 5C). Finally, extrinsic phycobilisomes (PCB) [34] were assembled on the irregular-arranged primary thylakoids (Fig.5D). So as a result, the inner cytoplasm with primary thylakoids looked like the cyanobacterial cytoplasm, but contained no cyanobacterial inclusions.

**Figure 5.**
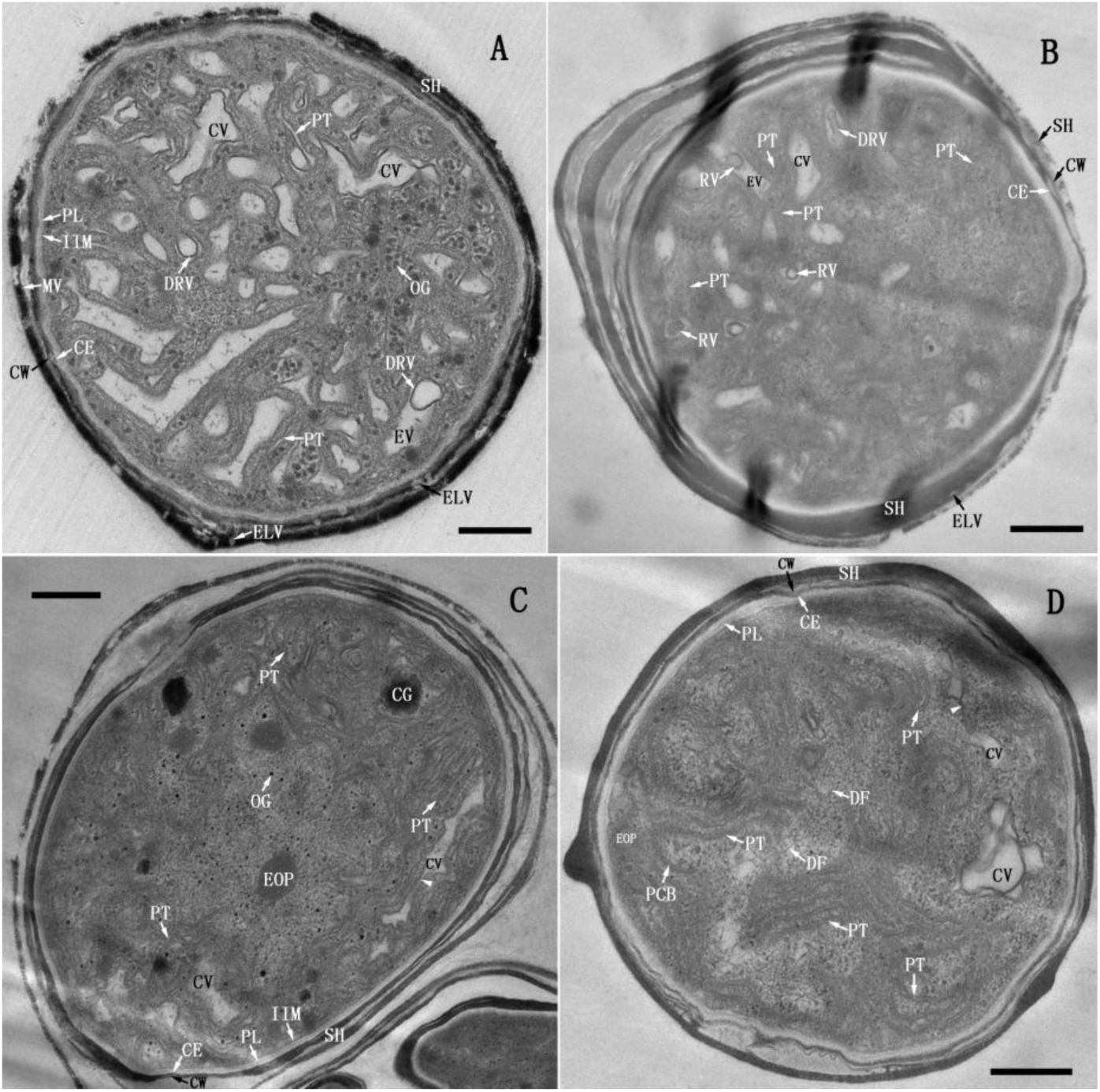
Formation of primary thylakoid (day 2-3). (**A**) Ring-shaped vesicles (RV) swelled up into dilated-ring-shaped vesicles (DRV), whose membranes met with the membranes of electron-transparent vesicles (EV), and thus gave rise to unit-membrane-bounded combined vesicles (CV). Subsequently, CV coalesced into longer ones or flattened out into slender short primary thylakoids (PT). (**B**) The newly formed short PT distributed randomly in the inner cytoplasm (ICP), whose matrix turned opaque. Occasionally a cluster of small RV presented in a EV. (**C**) The short PT extended or merged end-to-end into long PT; while the long CV flattened out into PT by localized-constriction (arrowhead). Meanwhile, several cyanophycin granules (CG) were formed. (**D**) The extrinsic phycobilisomes (PCB) were assembled on PT. Scale bar, 0.5 μm.

### De-compartmentalization, disassembly of primary thylakoids, DNA partition, re-compartmentalization and formation of secondary thylakoids

#### De-condensation and de-compartmentalization of inner cytoplasm, disassembly of primary thylakoids and DNA migration

As shown in Fig.6A, the inner cytoplasm de-condensed (solubilized) and became translucent with concomitant formation of less electron-dense materials (LDM), less electron-dense bodies (LDB) and cyanophycin granules; while the luminal matrix of primary thylakoids condensed, and so the membrane pair was in close apposition, seeming to be a single unit membrane with rough margin, between which short DNA fibers dispersed. In the meantime, the inner intracytoplasmic membrane disassembled into tiny vesicles (TV), such that the solubilized inner cytoplasm was de-compartmentalized and coalesced with the inner intracytoplasmic space; the less electron-dense materials diffused outward, blurring the compacted peptidoglycan-like layer, cytoplasmic envelope and cell wall (Fig. 6A). Subsequently, the coalesced inner cytoplasm and inner intracytoplasmic space was separated into lower and upper portions by the less electron-dense materials: in the lower portion, the primary thylakoids broke up into double-layered membrane fragments (DMF, two unit membranes) and thus DNA fibers aggregated; while in the upper portion, the double-layered membrane fragments began to curl and merge laterally into double-membraned vesicles (DMV) (Fig. 6B). As such, all double-layered membrane fragments were assembled into double-membraned vesicles, which dispersed along with DNA fibers in the coalesced inner cytoplasm and inner intracytoplasmic space (Fig. 6C). Thereafter, the double-membraned vesicles moved outward quickly and attached to the peptidoglycan-like layer that was cover by electron-dense materials, while the tangled DNA fibers migrated slowly resulting in an “empty” inner space (EIS), at the border of which the recruited tiny vesicles began to fuse and elongate into double-layered membrane segments (DMS) (Fig. 6D).

**Figure 6.**
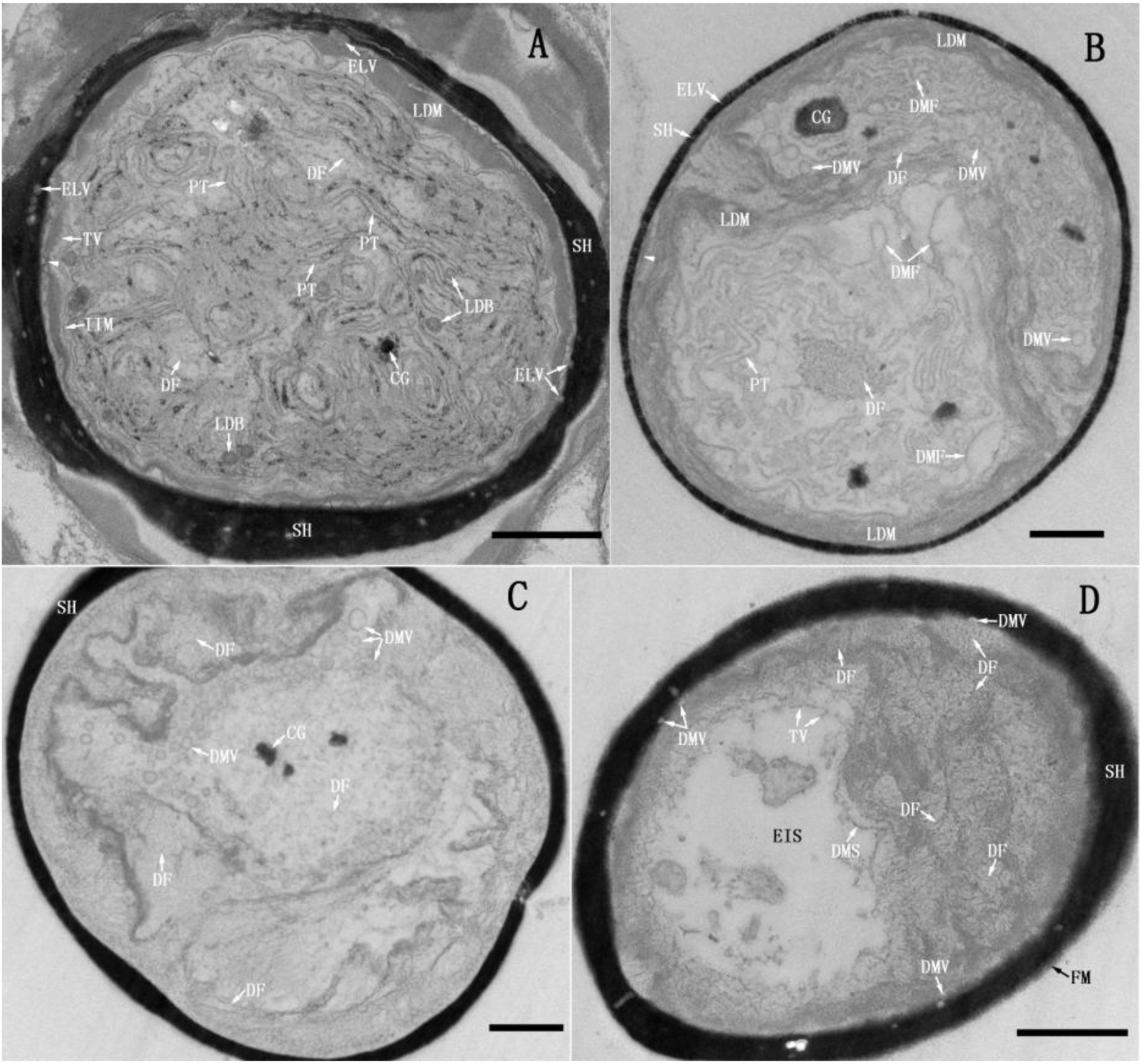
De-condensation and de-compartmentalization of inner cytoplasm, disassembly of primary thylakoids and DNA migration (day 4-5). (**A**) The inner cytoplasm (ICP) decondensed and became translucent, with concomitant production of less electron-dense materials (LDM), less electron-dense bodies (LDB) and cyanophycin granules (CG). The luminal matrix of primary thylakoids (PT) condensed and thus the membrane pair were in close apposition, between which short DNA fibers (DF) dispersed. The inner intracytoplasmic membrane (IIM) disassembled into tiny vesicles (TV), thus, the inner cytoplasm and inner intracytoplasmic space coalesced; LDM diffused outward blurring the compacted peptidoglycan-like layer (PL), cytoplasmic envelope (CE) and cell wall (CW) (arrowhead). (**B**) The primary thylakoids broke up into double-layered membrane fragments (DMF), which merged laterally into double-membraned vesicles (DMV); while DF aggregated into a cluster. (**C**) DMV and DF dispersed in the coalesced ICP and inner intracytoplasmic space (**D**) DMV moved outward quickly and attached to PL that was cover by electron-dense materials, while the tangled DF migrated slowly resulting in an “empty” inner space (EIS), at the border of which the recruited TV began to fuse and elongate into double-layered membrane segments (DMS). Scale bar, 0.5 μm.

#### Re-compartmentalization, DNA partition and formation of secondary thylakoids

As the double-layered membrane segments extended into the double-membraned intracytoplasmic envelope (ICE), the coalesced inner cytoplasm and inner intracytoplasmic space was re-compartmentalized into a new inner cytoplasm (NIC) and a new inner intracytoplasmic space (NIS) (Fig.7A). The new inner cytoplasm was enclosed by the intracytoplasmic envelope; while the new inner intracytoplasmic space represented the space between the intracytoplasmic envelope and peptidoglycan-like layer. Most DNA fibers were allocated into the new inner intracytoplasmic space, which de-condensed into cloudlike materials (CLM) or aggregated into thick DNA threads (DT); by contrast only few sporadic DNA fibers and electron-dense particles (EP) were partitioned into the new inner cytoplasm (Fig.7A). The intracytoplasmic envelope was not sealed, on the outer leaflet of which, some electron-transparent materials were synthesized, similar in appearance to the bacterial lipid [35]. Hereafter, accompanying the expansion of intracytoplasmic envelope, DNA fibers in the narrowing new inner intracytoplasmic space decondensed and likely attached to the thickened peptidoglycan-like layer (Fig.7B); while the double-membraned vesicles that were covered by less electron-dense materials detached from the peptidoglycan-like layer, moving inward via the opening of intracytoplasmic envelope into the new inner cytoplasm and outward through the pores on the peptidoglycan-like layer into the outer intracytoplasmic space (Fig.7B). Hence, the fenestrated peptidoglycan-like layer served not only as a mechanical and osmotic barrier, but also a platform for anchoring DNA and double-membraned vesicles. When the intracytoplasmic envelope closed up, DNA in the new inner intracytoplasmic space was assembled into thick DNA fibers resembling chromatin fibers [36] with concomitant formation of countless ribosomes (Fig. 7C). Simultaneously, in the new inner cytoplasm an increased number of DNA fibers and many ribosomes were synthesized; the double-membraned vesicles dilated, opened up and became double-layered membrane fragments (Fig. 7C). After that, the double-layered membrane fragments extended randomly into spiral thylakoids, which were devoid of phycobilisomes and morphologically different from the primary thylakoids, termed secondary thylakoids (ST) (Fig.7D). Concomitant with development of secondary thylakoids was the formation of osmiophilic granules and electron-opaque bodies as well as enrichment of DNA fibers and ribosomes in the expanded new inner cytoplasm (Fig.7D). The structures outside of the new inner cytoplasm were fuzzy owing to diffusion of electron-dense materials. The major portion of peptidoglycan-like layer was dismantled, and thus the new inner intracytoplasmic space and outer intracytoplasmic space coalesced into a new intracytoplasmic space (NS), whose content was termed intracytoplasmic matrix (NX). There were different vesicles in the new intracytoplasmic space, including double-membraned vesicles as well as newly formed electron-translucent oblong vesicles (OV) and electron-opaque vesicles (EOV) (Fig.7D).

**Figure 7.**
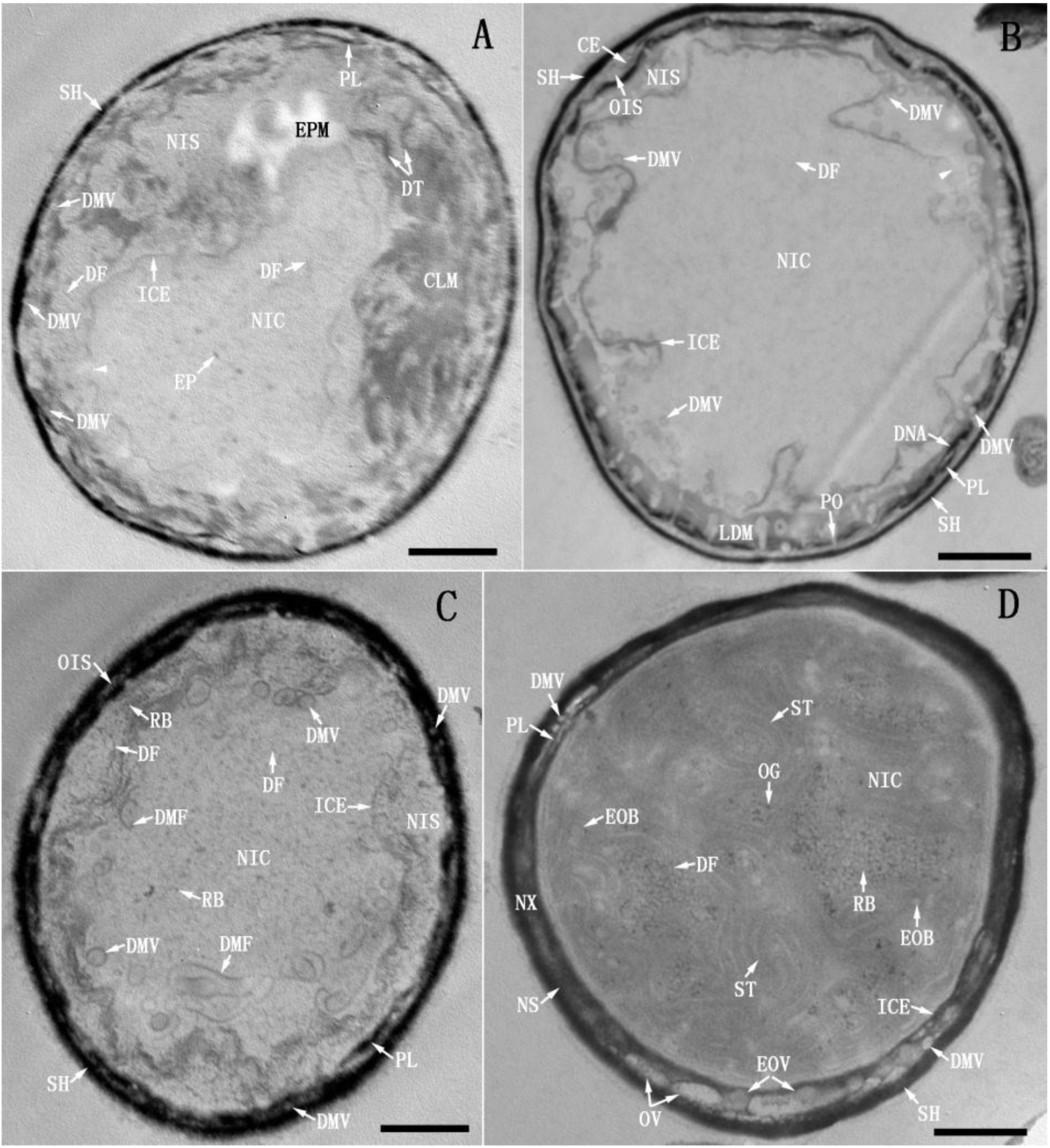
Re-compartmentalization, DNA partition and formation of secondary thylakoids. (day 5-6) (**A**) The double-membraned intracytoplasmic envelope (ICE) re-compartmentalized the coalesced inner cytoplasm and inner intracytoplasmic space into a new inner cytoplasm (NIC) and a new inner intracytoplasmic space (NIS). Most of the DNA fibers (DF) were allocated into NIS, which decondensed into cloudlike materials (CLM) or aggregated into DNA threads (DT), while only few DF and electron-dense particles (EP) were partitioned into NIC. ICE had an opening (arrowhead), on the outer leaflet of which, some electron-transparent materials (EPM) were synthesized. (**B**)The double-membraned vesicles (DMV) in NIS moved into NIC via ICE opening (arrowhead), or passed through the pores (PO) on the peptidoglycan-like layer (PL) into outer intracytoplasmic space (OIS). (**C**) ICE sealed, DNA in NIS condensed into a mass of DF with concomitant formation of countless ribosomes (RB); while an increased number of DF and RB were formed in NIC; DMV opened up into double-layered membrane fragments (DMF) and elongated. (**D**) DMF in NIC extended into secondary thylakoids (ST) with concomitant formation of osmiophilic granules (OG) and electron-opaque bodies (EOB) as well as enrichment of DF and RB. Outside of NIC, the major portion of PL was dismantled, such that NIS and OIS coalesced into a new intracytoplasmic space (NS), sequestering new intracytoplasmic matrix (NX), DMV, oblong vesicles (OV) and electron-opaque vesicles (EOV). Scale bar, 0.5 μm.

The viability of TDX16 manifested the coordination and competence of the only two compartments in performing cellular functions. The new inner cytoplasm performed photosynthesis and respiration on the secondary thylakoids just like the cases of cyanobacteria, though it contained only a handful of DNA and no cyanobacterial inclusion except osmiophilic granules; while the new intracytoplasmic matrix retained the major fraction of cellular DNA and performed most if not all of other metabolic activities.

### Biogenesis of primitive chloroplast, eukaryotic cell wall and primitive nucleus

#### Biogenesis of primitive chloroplast and eukaryotic cell wall

As shown in Fig.8A, the new inner cytoplasm became polarized, in which the newly formed secondary thylakoids underwent disassembly, leaving some remnants in the lower region; while parallel arrays of discrete slender sacs with transparent matrix were being developed in the upper region. These parallel-arranged slender sacs were morphologically similar to algal and plant thylakoids, termed primitive eukaryotic thylakoids (PMT), which appeared to develop from the plastoglobuli (similar in appearance to but smaller in size than osmiophilic granules) formed during disassembly of secondary thylakoids (Fig.8A), in a way similar to development of primary thylakoids from osmiophilic granules (Fig.5). Beneath the primitive eukaryotic thylakoids, a nascent pyrenoid (PD) with an incomplete starch plate (SP) and two starch granules (SG) were formed (Fig. 8A), both of which were the characteristic bodies of green algal chloroplasts [37]. Accordingly, the new inner cytoplasm developed into a primitive chloroplast (PC) delimited by the double-membraned intracytoplasmic envelope. That was to say, the intracytoplasmic envelope became the double-membraned chloroplast envelope (CHE). The absence of mitochondrion suggested that respiration took place on the primitive eukaryotic thylakoids. Thus, the primitive chloroplast was a dual-function composite organelle.

**Figure 8.**
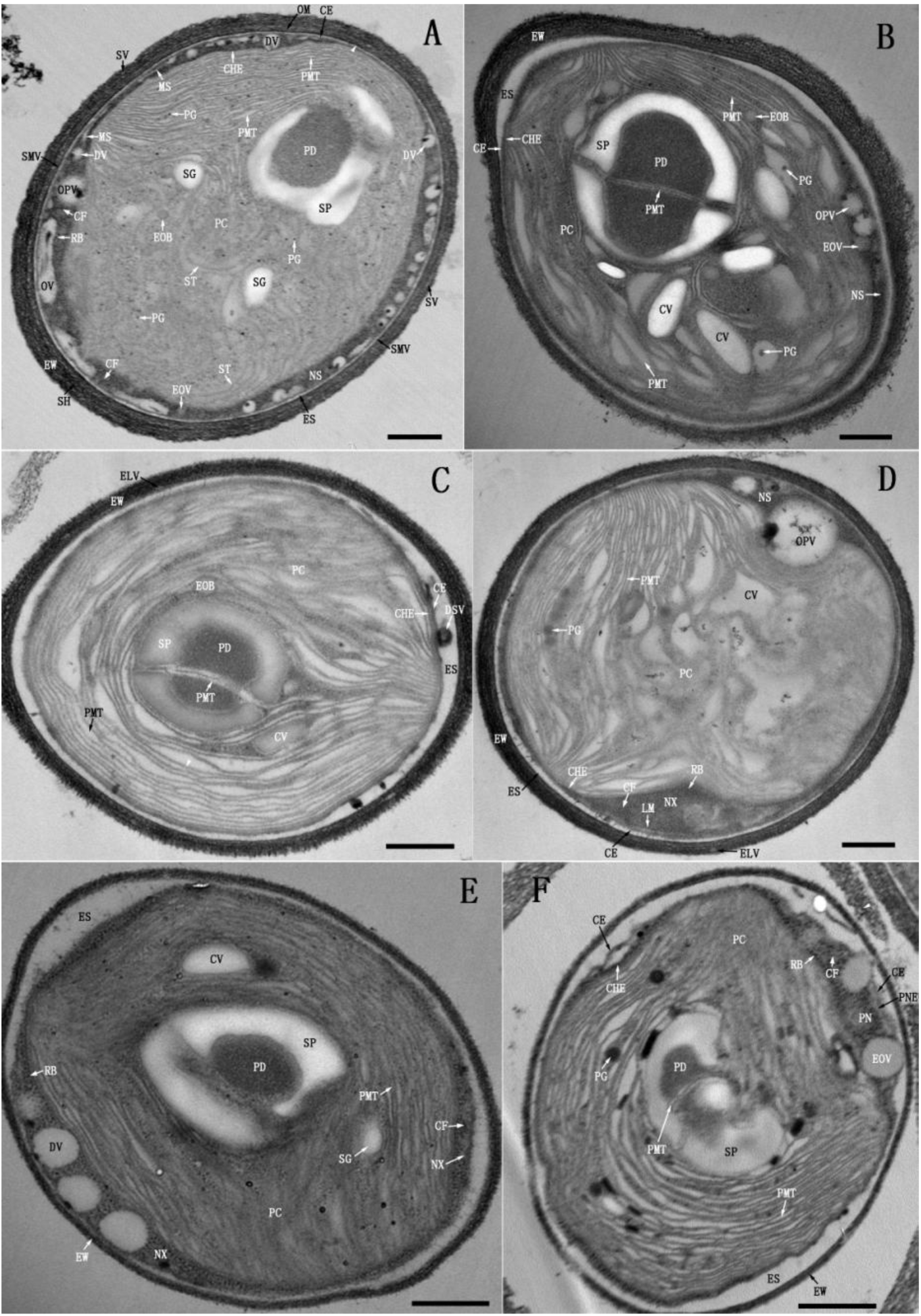
Biogenesis of primitive chloroplast, eukaryotic cell wall and primitive nucleus (day-7-9). (**A**) The secondary thylakoids (ST) were dismantled leaving some remnants and plastoglobuli (PG) in the lower region of new inner cytoplasm (NIC), while parallel arrays of primitive eukaryotic thylakoids (PMT) were being developed in the upper region of NIC with concomitant formation of a nascent pyrenoid (PD) with an incomplete starch plate (SP), and two starch granules (SG). So, NIC developed into the primitive chloroplast (PC), and the intracytoplasmic envelope became the chloroplast envelope (CHE). Inside the new intracytoplasmic space (NS), thick chromatin fibers (CF) and large ribosomes (RB) were formed; the peptidoglycan-like layer and the double-membraned vesicles disappeared; while many small dotted vesicles (DV) and opaque-periphery vesicle (OPV) emerged. Some DV began to fuse and flattened into membrane segments (MS). Outside of the new intracytoplasmic space (NS), some smaller vesicles (SMV) shed from the cytoplasmic envelope (CE) into the extracytoplasmic space (ES); the peptidoglycan layer (P) turned into an electron-dense layer (EL) (arrowhead), and a stratified SH embedded with small vesicles (SV) was formed external to the outer membrane (OM). Hence OM, EL and the sandwiched electron-transparent space constituted a trilaminar domain, which along with SH became the eukaryotic cell wall (EW). (**B**) PMT with wide luminal space were formed continuously by elongation of the combined vesicle (CV); PD was surrounded with a complete SP and bisected by two pairs of PMT; hence PC expanded and occupied most NS in the longitudinally sectioned plane; (**C**) PC filling with PMT occupied whole NS in longitudinally sectioned plane; CHE adhered to CE and a dense vesicle (DSV) shed off from CE into the widened ES. (**D**) The vertical section of a cell. The anterior portion of PC contacted CE; MS merged into the limiting membrane (LM) at the border of new intracytoplasmic matrix (NX) (**E**) The oblique section of a cell. The major fraction of NX converged at one side of PC. (**F**) The tangential section of a cell. NX was encased by LM into the primitive nucleus (PN) containing CF, RB and electron-opaque vesicles (EOV); while LM became primitive nuclear envelope (PNE). NS vanished, and so CE shrank and wrapped PC and PN. Scale bar, 0.5 μm.

The new intracytoplasmic space became clear: the new intracytoplasmic matrix condensed, containing ribosomes and chromatin fibers (CF) [36, 38]; the peptidoglycan-like layer and double-membraned vesicles disappeared, while many small dotted vesicles (DV) blistered from the chloroplast envelope and lined up along the cytoplasmic envelope, some of which began to fuse and flattened out into membrane segments (MS) (Fig.8A). Such a scenario of membrane synthesis was akin to nuclear envelope assembly [39–40]. In addition, a large coated-vesicle-like opaque-periphery vesicle (OPV) was being assembled at the primitive chloroplast envelope, which bridge the primitive chloroplast and cytoplasmic envelope probably for transferring substances.

The fuzzy electron-dense sheath external to the cell wall (Fig.7D) scaled off, while a stratified sheath embedded with many small vesicles formed, which adhered to the outer membrane and made the latter difficult to discern (Fig.8A); the peptidoglycan layer (Fig. 4A) became denser and thicker indicating changes of its composition and thus was referred to as electron-dense layer (EL). In this case, the outer membrane along with the electron-dense layer and the sandwiched electron-transparent space still adopted its original configuration (Fig. 4A), and thus constituted a trilaminar domain resembling the trilaminar sheath in the cell walls of green algae [41–45], which combined with the stratified sheath into a continuum similar in structure to the cell walls of green algae, termed eukaryotic cell wall (EW).

#### Biogenesis of primitive nucleus

As plastoglobuli developed progressively into combined vesicles and then flattened into primitive eukaryotic thylakoids with wide luminal space in the lower region of primitive chloroplast, the pyrenoid got matured, which was surrounded with a complete starch plate and bisected by two pairs of primitive eukaryotic thylakoids. Hence, the primitive chloroplast expanded substantially with corresponding shrinkage of the new intracytoplasmic space (Fig. 8B). Subsequently, the primitive eukaryotic thylakoids coalesced and extended around the pyrenoid, the adjacent membranes of which were connected, seeming to be a single membrane (Fig. 8C); the further expanded primitive chloroplast occupied whole new intracytoplasmic space in the longitudinally sectioned planes, and so the primitive chloroplast envelope fully adhered to the cytoplasmic envelope, from the latter of which a dense vesicle (DSV) shed off into the widened extracytoplasmic space (Fig.8C). Consistently, vertical profile (Fig.8D) showed that the anterior portion of new intracytoplasmic space disappeared owing to expansion of primitive chloroplast, as such the new intracytoplasmic matrix in the shrunken new intracytoplasmic space was concentrated by squeezing out liquid into the extracytoplasmic space, at the border of which the membrane segments coalesced into a limiting membrane (LM) (Fig.8D). As the expansion of primitive chloroplast continued, the stepwise-concentrated new intracytoplasmic matrix moved to (Fig. 8E) and finally converged at one side of primitive chloroplast, which was ensheathed by the limiting membrane into a membrane-bounded organelle adhering to the primitive chloroplast (Fig. 8F). This organelle sequestered the new intracytoplasmic matrix that retained the major fraction of cellular DNA and performed metabolic activities, and thus was also a composite organelle, termed primitive nucleus (PN). As such, the limiting membrane became the primitive nucleus envelope (PNE) that probably consisted of four unit membranes, because it was synthesized by fusion of the small dotted vesicles that were budded from the primitive chloroplast envelope (Fig. 8A) and likely delimited by two unit membranes. Concurrent with the formation of primitive nucleus, the new intracytoplasmic space vanished, and thus the cytoplasmic envelope shrank and wrapped the only two primitive organelles, apparently to keep intercommunication via intimate membrane contacts in this vital transient state (Fig. 8F).

### Formation of eukaryotic cytoplasmic matrix and biogenesis of mitochondria

#### Concurrent formation of eukaryotic cytoplasmic matrix and biogenesis of mitochondria

As shown in Fig.9A, a vesicle-containing body (VB), apparently chipped off the invaginated primitive chloroplast, was being engulfed by the primitive nucleus with concomitant formation of a thin layer of electron-dense materials; and concurrently a small nascent mitochondrion (M) was being assembled within the primitive chloroplast. The primitive nucleus envelope was contiguous with the cytoplasmic envelope at the outer side, but separated into two sets of double-membraned envelopes inside the primitive chloroplast cavity, the inner and outer sets of which were referred to as nuclear envelope (NE) and outer nuclear envelope (OE) respectively. This result confirmed that the primitive nucleus envelope consisted of four unit membranes. The thin layer of electron-dense materials was a ribosome-containing fluid extruded from the primitive nucleus at the site where the nuclear envelope and outer nuclear envelope fused, which served as the medium connecting the primitive organelles and cytoplasmic envelope, and thus was the incipient eukaryotic cytoplasmic matrix (cytosol) (EM). After ‘digestion’ of the vesicle-containing body, the primitive nucleus and eukaryotic cytoplasmic matrix both increased in sizes; while an oval mitochondrion with characteristic cristae (CR) emerged in the apical dome of the enlarged primitive chloroplast cavity (Fig.9B). The nuclear envelope and outer nuclear envelope were separated by an interenvelope space (IES), but merged at one site resulting in a wide opening, from which the primitive nuclear matrix was extruded. Ribosomes in the eukaryotic cytoplasmic matrix were larger than those within the primitive chloroplast stroma (SM), most of which were bound to the outer membranes of primitive organelles (Fig. 9B). During this process, a number of smaller vesicles and microfibrils (ML) budded and emanated from cytoplasmic envelope into extracytoplasmic space, respectively (Fig. 9B). Concurrent extrusion of primitive nuclear matrix and formation of mitochondrion in different cells took place in the same manner (Fig. 9C, D and E), and in one of which two spindle-shaped mitochondria were just being assembled within the primitive chloroplast (Fig. 9D). Occasionally, an unusual membrane-delimited intranuclear body (IB) appeared in the primitive nucleus (Fig. 9F), which seemed to be developed during engulfment and digestion of the vesicle-containing body or the like and play a role in selective extrusion of the primitive nuclear matrix to build up the eukaryotic cytoplasmic matrix.

**Figure 9.**
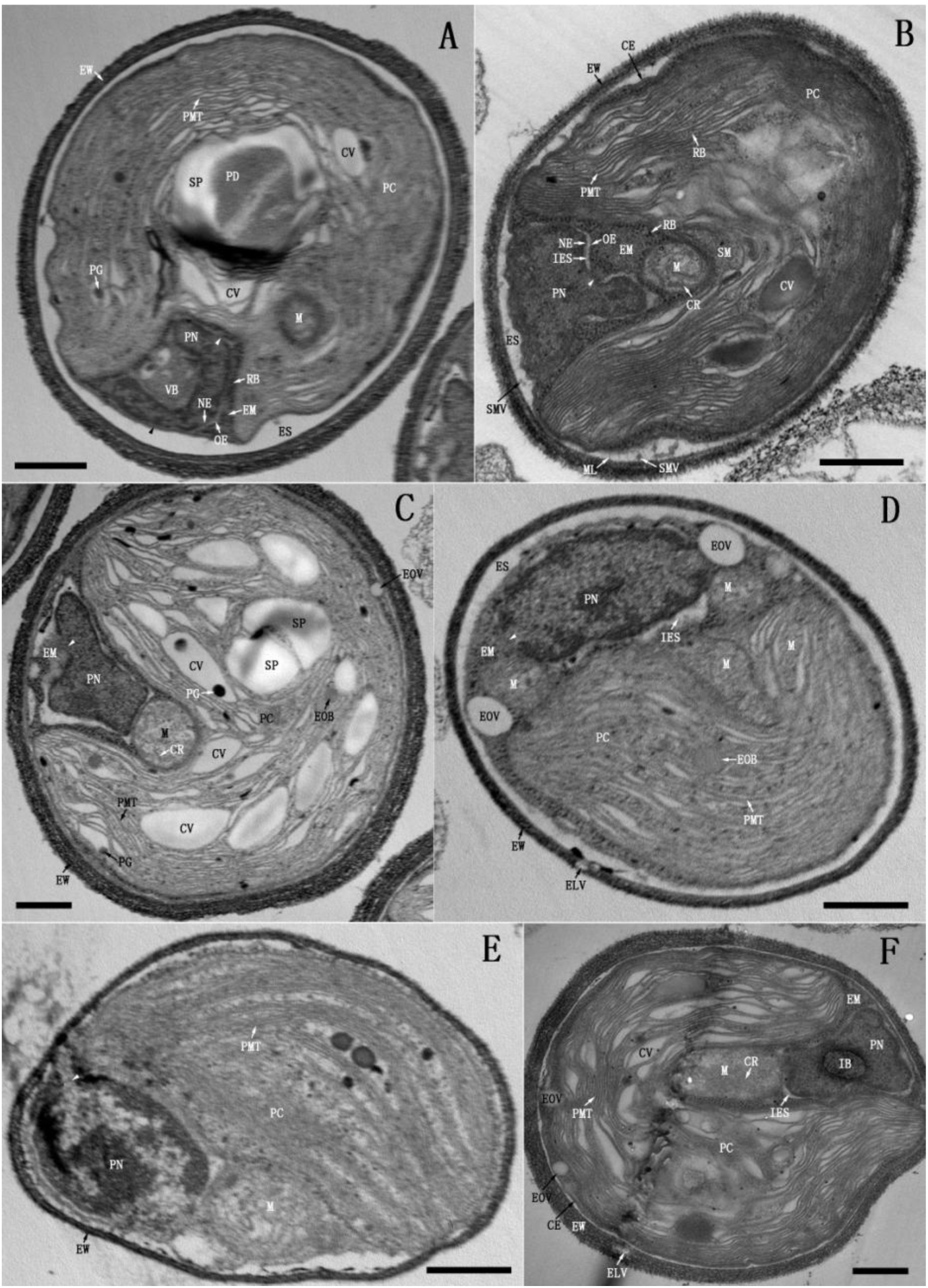
Concurrent formation of eukaryotic cytoplasmic matrix and biogenesis of mitochondria (day 8-9). (**A**) The primitive nucleus (PN) engulfed a vesicle-containing body (VB) and extruded nuclear matrix (white arrowhead) to build up the eukaryotic cytoplasmic matrix (EM); while a mitochondrion (M) was assembled in the primitive chloroplast (PC). The nuclear envelope (NE) and outer nuclear envelope (OE) were separated in PC cavity but contiguous with the cytoplasmic envelope (CE) at the outer side (black arrowhead). (**B**) PC with enriched stroma (SM) further invaginated, in the apical dome of its cavity, a mitochondrion with characteristic cristae (CR) emerged. OE and NE were separated by an inter envelope space (IES), but fused at one site into a large opening (arrowhead), from which the nuclear matrix was extruded. A number of smaller vesicles (SMV) and microfibrils (ML) budded and emanated respectively from CE into the extracytoplasmic space (ES). (**C**), (**D**) and (**E**) PN extruded nuclear matrix (arrowhead). (**F**) PN contained a membrane-delimited intranuclear body (IB). Scale bar, 0.5 μm.

#### Continuous biogenesis of mitochondria after formation of eukaryotic cytoplasmic matrix

After building up sufficient eukaryotic cytoplasmic matrix, bulk extrusion of primitive nuclear matrix ceased. So the primitive nucleus got matured into a nucleus (N) and TDX16 developed into a premature eukaryote, because new mitochondria were continuously developed in the primitive chloroplast even after the formation of vacuole (V) with internal vesicle (IV), multilamellar body (MLB), lipid droplet (LD), and small opaque vesicle (SOV), leading to distortion of the primitive eukaryotic thylakoids and diminishment of the primitive chloroplast (Fig.10). As shown in Fig.10A, a small mitochondrion was being developed in the periphery of the primitive chloroplast; while a twisting dumbbell-shaped mitochondrion in another cell was nearly finished, one of its bulbous-end sequestering an internal body (ITB) was segregated, but another end still under development in the primitive chloroplast (Fig.10B). Details of mitochondrion biogenesis were detected during assembly of giant mitochondria. As shown in Fig. 10C, a giant ‘L-shaped’ mitochondrion was in development and still continuous with the primitive chloroplast in the region around its corner point: the inner side envelope of its long arm and the corresponding portion of chloroplast envelope, as well as the interior cristae were nearly complete; while those of its short arm were just being synthesized. All these membranes were synthesized by merging the small dense-margined vesicles (DGV) developed from the segmented primitive eukaryotic thylakoids (Fig. 10C). Similarly, in another cell a bulky mitochondrion was undergoing segregation, which was connected with the primitive chloroplast on the inner side but a small mitochondrion on the outer side (Fig.10D). In the inner and outer interfaces, three and two pairs of contorted membranes were being synthesized respectively by fusion of dense-margined vesicles: among the three pairs of membranes, the outer and middle ones were the segments of mitochondria envelope and chloroplast envelope respectively, while the inner one was likely the envelope of the next mitochondrion that appeared to be in preparation; likewise the two pairs of membranes were the outer side and inner side mitochondria envelope of the bulky and small mitochondria respectively (Fig.10D).

**Figure 10.**
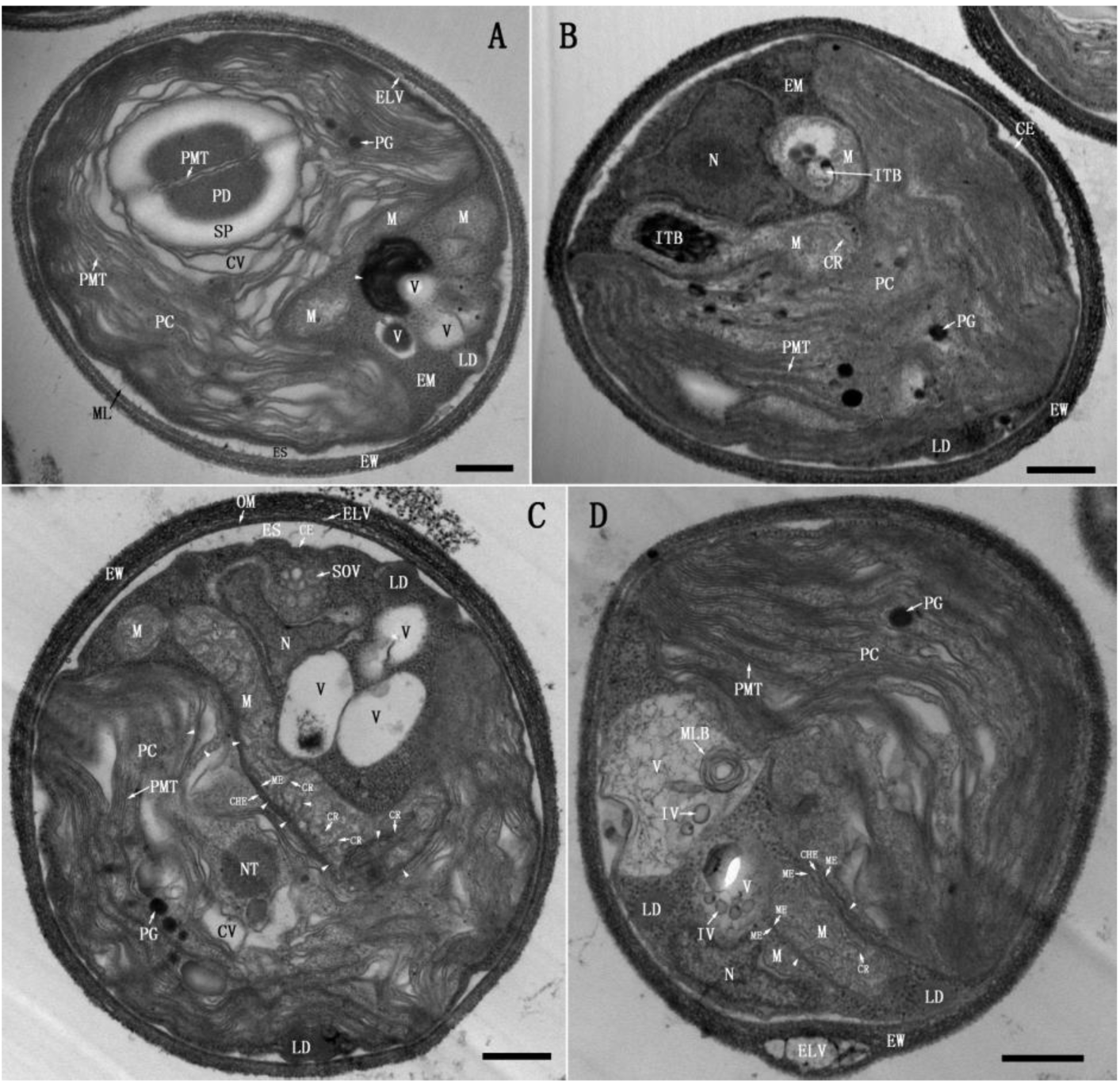
Biogenesis of mitochondria after building up eukaryotic cytoplasmic matrix (day 8-9). (**A)** A small mitochondrion was being developed in the primitive chloroplast (PC) in the presence of two mitochondria, three vacuoles (V), a lipid droplet (LD) and some electron-dense materials (arrowhead). (**B**) A twisting dumbbell-shaped mitochondrion was nearly finished, one of its bulbous-end sequestering an internal body (ITB) was segregated, but the other end was contiguous with PC (**C**) A large ‘L-shaped’ mitochondrion was being assembled in the presence of three vacuoles and a cluster of small opaque vesicles (SOV), which was continuous with PC in the region around its corner point. The inner side mitochondrial envelope (ME) and the corresponding portion of chloroplast envelope (CHE) as well as cristae (CR) were being synthesized by fusion of the dense-margined vesicles (DGV) (arrowhead) that were developed by segmentation of the primitive eukaryotic thylakoids (PMT). There was a large nucleoid-like structure (NT) in the venter side of PC. (**D**) After emergence of the large vacuole with internal vesicles (IV) and a multilamellar body (MLB), a bulky mitochondrion was being developed, which was connected with PC on the inner side but a small mitochondrion on the outer side. In the inner and outer interfaces, three and two pairs of contorted membranes were being synthesized respectively by merging DGV (arrowhead). In addition, several large electron-translucent vesicles (ELV) were embedded in the eukaryotic cell wall (EW). Scale bar, 0.5 μm.

The above results indicated that mitochondria were assembled in the periphery of primitive chloroplast by encapsulating selected components with the membranes derived from the primitive eukaryotic thylakoids. As the assembly nearly finished, the chloroplast envelope opened up at the ventral side allowing mitochondria to detach by twisting, and then resealed by incorporating the membrane segment concurrently synthesized with the mitochondrial envelope. Since mitochondria were always assembled in the ventral side of primitive chloroplast where a large nucleoid-like structure situated (Fig.10C), it seemed likely that DNA was synthesized in the nucleoid-like structure and subsequently sorted into the mitochondria.

### Transition of mitochondria into vacuoles and degradation of primitive eukaryotic thylakoids-derived vesicles

#### Transition of mitochondria into double-membraned vacuoles

Following the emergence of new mitochondrion, the opaque matrix of previously formed mitochondria began to self-degrade into electron-transparent material (Fig.11A), such that the mitochondria turned into double-membraned vacuoles (V) with electron-transparent matrix, containing incompletely degraded internal body, vesicle and cristae (Fig. 11B). Most of the residual inclusions were further degraded into electron-dense debris (ED) (Fig.11C) and released into the eukaryotic cytoplasmic matrix (Fig. 11D); while some internal bodies themselves also contained small internal bodies, so after degradation of their contents, the residual membranes (Fig. 11D) were collected into multilamellar bodies [46–47] (Fig, 11 E and F). Accompanying the development of vacuoles, a new mitochondrion was assembled at the edge of primitive chloroplast (Fig.11B); a piece of chloroplast debris (CD) emerged in the eukaryotic cytoplasmic matrix (Fig.11B); and lipid droplets formed intensively on the inner membrane of cytoplasmic envelope and the outer membrane of primitive chloroplast envelope (Fig.11B,C,D,E,F).

**Figure 11.**
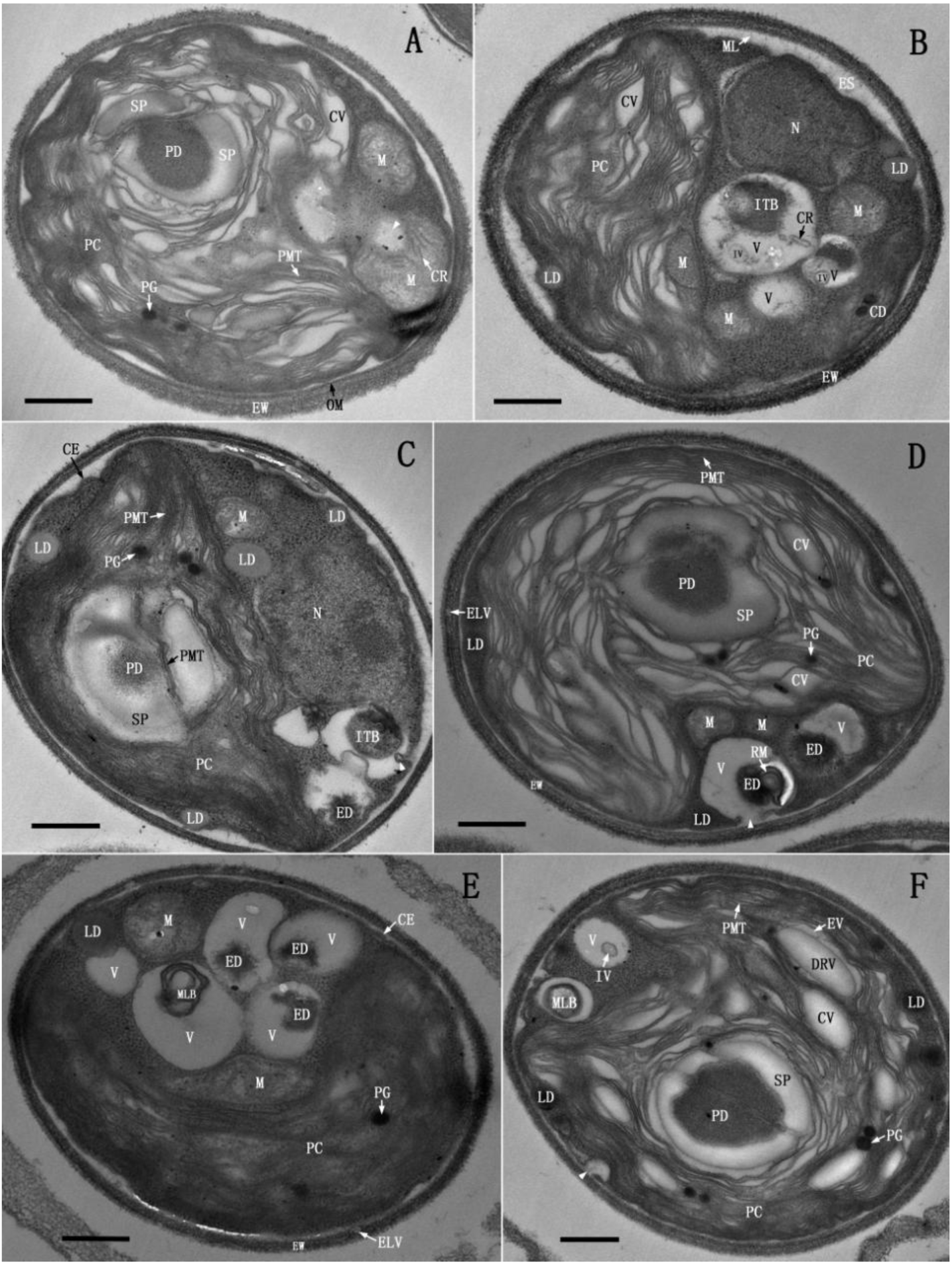
Transition of mitochondria into double-membraned vacuoles (day 8-9). (**A**) The matrix of a mitochondrion was being degraded (arrowhead). (**B**) After matrix degradation, the mitochondria turned into vacuoles (V) containing internal body (ITB), internal vesicle (IV) and remnant cristae (CR) or only electron-transparent matrix. Meanwhile, a new mitochondrion was developed in the primitive chloroplast (PC), a lipid droplet (LD) was formed at the inner leaflet of cytoplasmic envelope (CE) and a piece of chloroplast debris (CD) emerged in the eukaryotic cytoplasmic matrix (EM). (**C**) ITB in two small vacuoles degraded into electron-dense debris (ED); while ITB in the large vacuole remained intact, the vacuolar membranes fused with CE and invaginated (arrowhead). (**D**) a small vacuole expelled ED into EM; while a large vacuole sequestered some residual membranes (RM), the membranes of which fused with CE, resulting in an opening (arrowhead). (**E**) A large vacuole contained a multilamellar body (MLB). (**F**) Two vacuoles contained a MLB and an IV respectively. A small vacuole fused with CE, resulting in an opening (arrowhead); while several dilated ring-shaped vesicles (DRV) emerged in electron-transparent vesicles (EV). Scale bar, 0.5 μm.

#### Formation and degradation of primitive eukaryotic thylakoids-derived vesicles and coalescence of vacuoles

After vacuoles came into being, the short fragmented primitive eukaryotic thylakoids that were generated during assembly of mitochondria curled and ‘rolled up’ into “vesicle within vesicle” like compound vesicles (CPV) (Fig.12 A,B and C), which were internalized by vacuoles directly as they segregated from the primitive chloroplast (Fig. 12C), or after they shed into the eukaryotic cytoplasmic matrix (Fig.12A and B). And then contents of the internalized compound vesicles were degraded, while the remaining membranes stacked up into multilamellar bodies within the vacuoles (Fig. 12D, E and F), which began to fuse with each other by membrane protrusion. As shown in Fig. 12C and D, a vacuole protruded into another one and fused at the contact sites, such that the membrane protrusion pinched off and became an internal vesicle (Fig. 12E) or shed as membrane fragments (Fig. 12 D and F). When the primitive chloroplast dwindled to a normal size, no more mitochondria and compound vesicles were produced and all vacuoles coalesced into a large one (Fig.12F). Accordingly, the primitive chloroplast got matured into chloroplast (C), and the primitive eukaryotic thylakoids matured into eukaryotic thylakoids (T).

**Figure 12.**
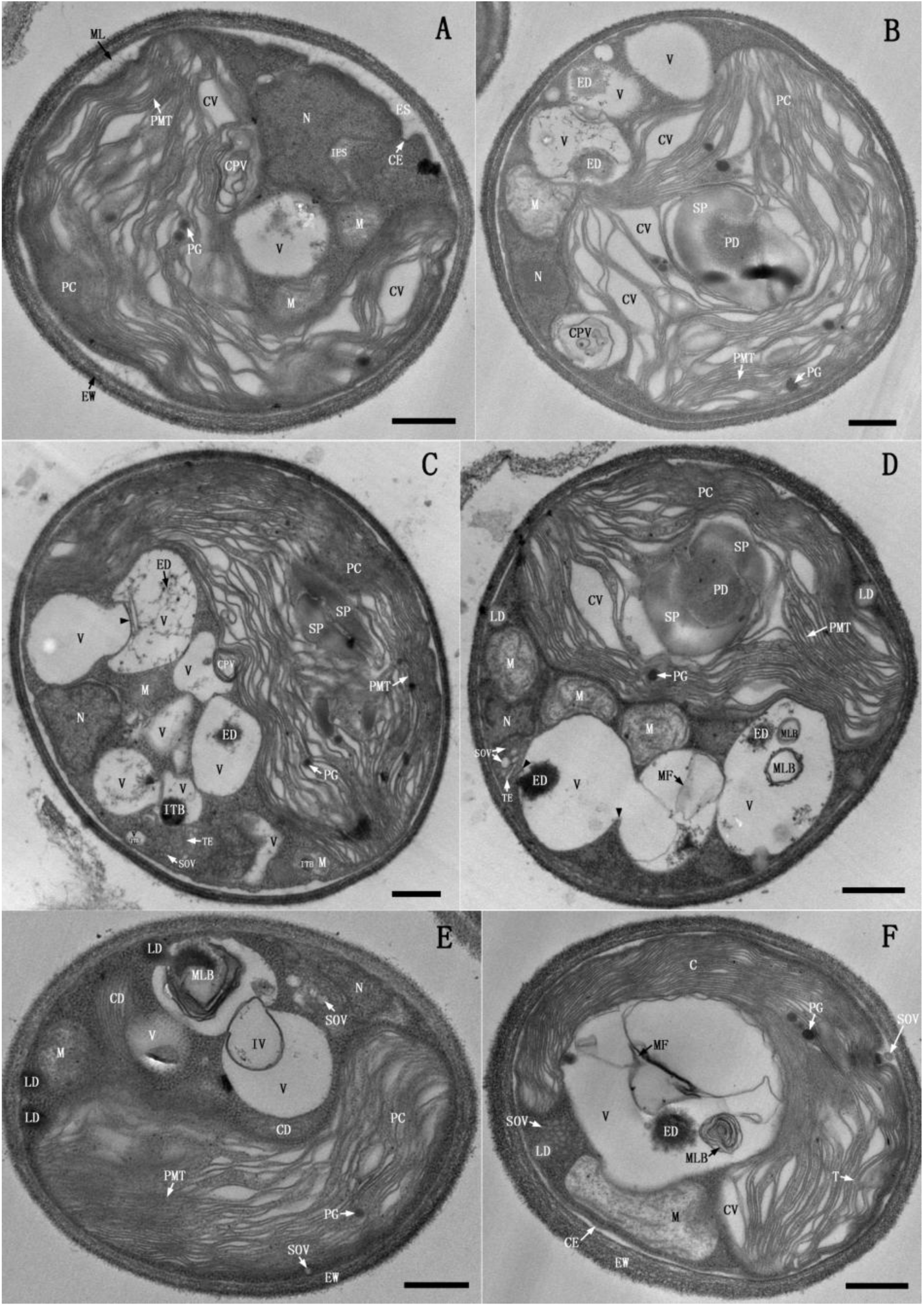
Formation and degradation of primitive eukaryotic thylakoids-derived vesicles and coalescence of vacuoles (day 8-9). (**A**) A collection of short primitive eukaryotic thylakoids (PMT) ‘rolled up’ into “vesicle within vesicle” like compound vesicle (CPV) in the margin of primitive chloroplast (PC). (**B**) A CPV was segregating from PC into the eukaryotic cytoplasmic matrix (EM). (**C**) A CPV was detaching from PC into a vacuole; while a large vacuole protruded into another one (arrowhead). (**D**) Membranes of the protruded vacuole and the vacuole that contained membranous fragments (MF) fused at their contact site (arrowhead). (**E**) Two vacuoles were merging; while a conspicuous piece of chloroplast debris (CD) presented in EM. (**F**) PC matured into a chloroplast (C); all vacuoles coalesced into a single vacuole containing electron-dense debris (ED), MF and MLB. Scale bar, 0.5 μm.

### Vacuole mediated unconventional exocytosis and endocytosis

#### Vacuole-mediated unconventional exocytosis

Vacuoles came into contact with the cytoplasmic envelope, and then the contacted membranes fused or broke up into fragments resulting in openings, from which a small quantity (Fig. 11D, F and Fig. 13A) or a large amount of vacuolar contents and small opaque vesicles that seemed to be internalized from the neighboring vesicle clusters were expelled into the extracytoplasmic space (Fig.13B). Apparently, vacuole-mediated exocytosis was unconventional exocytosis [48–50], as the exocytotic proteins were not sorted through Golgi apparatus.

**Figure 13.**
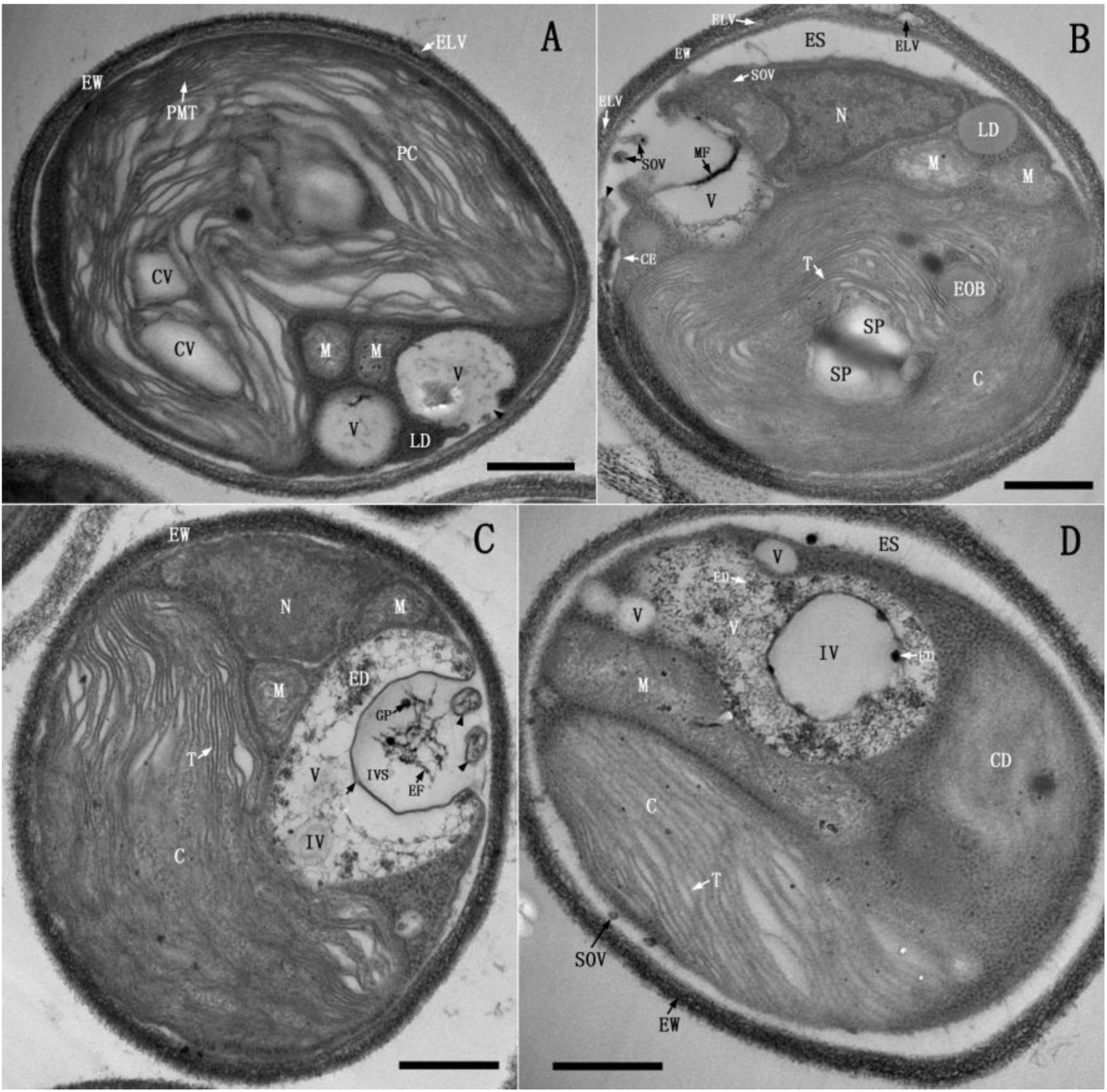
Vacuole mediated unconventional exocytosis and endocytosis (day 8-9). (**A)** An opening was formed at the contact site of the vacuolar membranes and cytoplasmic envelope (CE) (arrowhead), from which the vacuolar content was released. (**B**) The contacted vacuolar membranes and CE broke up into fragments (arrowhead), resulting in a wide opening, from which the soluble contents and small opaque vesicle (SOV) were expelled outside into the extracytoplasmic space (ES). (**C**) The vacuolar membranes merged with CE at two distant sites and then invaginated, resulting in a large invaginated space (IVS) entrapping some electron-dense fibrils (FB) and globular particles (GP); while CE between the two merged sites disrupted and coiled into membranous structures (arrowhead). It was clear that the vacuolar membranes consisted of two unit membranes (arrow). (**D**) A vacuole contained a large internal vesicle (IV). Scale bar, 0.5 μm.

#### Vacuole-mediated unconventional endocytosis

The vacuolar membranes merged with cytoplasmic envelope at two distant sites and then invaginated, resulting in a large invaginated space (IVS), entrapping some electron-dense fibrils (EF) and globular particles (GP); while cytoplasmic envelope between the two merged sites disrupted and coiled into membranous structures (Fig.13C). Upon the invaginated space reaching a certain size, the membrane invagination pinched off into the vacuole lumen and became a large internal vesicle (vacuole?), whose content was degraded in situ or discharged into the vacuole lumen (Fig.13D). In the same way, the nascent vacuoles also mediated small episode of endocytosis (Fig.10D and Fig.11C). Evidently, vacuole-mediated endocytosis was unconventional endocytosis, as there was no other organelle involved.

### Transition of TDX16 into an eukaryotic alga TDX16-DE

After bulk exocytosis and endocytosis, the large vacuoles vanished and no other organelle was formed. Thus, the prokaryotic cyanobacterium TDX16 (Fig.2-3) turned into a stable eukaryotic alga TDX16-DE with unique structure (Fig.14A, B and C). TDX16-DE cell has an eukaryotic cell wall, an extracytoplasmic space and a double-membraned cytoplasmic envelope, contains a chloroplast, a nucleus with two set of envelopes, two mitochondria, and no or several double-membraned vacuoles, but lacks endoplasmic reticulum, Golgi apparatus and peroxisome.

The nucleus still retains ribosomes (Fig.14A, B and C), indicating its capability of protein synthesis. The nuclear envelope and outer nuclear envelope have no visible pores and usually connect with the cytoplasmic envelope at the opening of chloroplast cavity (Fig.14A). When they are separated, a number of electron-dense vesicles (EDV) bud form the nuclear envelope into the interenvelope space, which fuse with and re-bud from the outer nuclear envelope and ultimately migrate to the two sides of chloroplast envelope and cytoplasmic envelope (Fig.14B), probably transferring nucleus-synthesized proteins for membrane renewal. Meanwhile, several openings are formed on the outer nuclear envelope, one of which is at the site where it merges with the cytoplasmic envelope. Such that, the eukaryotic cytoplasmic matrix and extracytoplasmic space is connected through the interenvelope space, enabling the exchange of metabolites between these two compartments (e.g., protein secretion) (Fig. 14B). By contrast, when the outer nuclear envelope and nuclear envelope come into contact, they disrupt at several sites and result in openings, allowing nucleocytoplasmic transport (Fig.14C), and fuse with the cytoplasmic envelope at one site resulting in a fusion pore, enabling the direct communication between nucleus and extracytoplasmic space.

**Figure 14.**
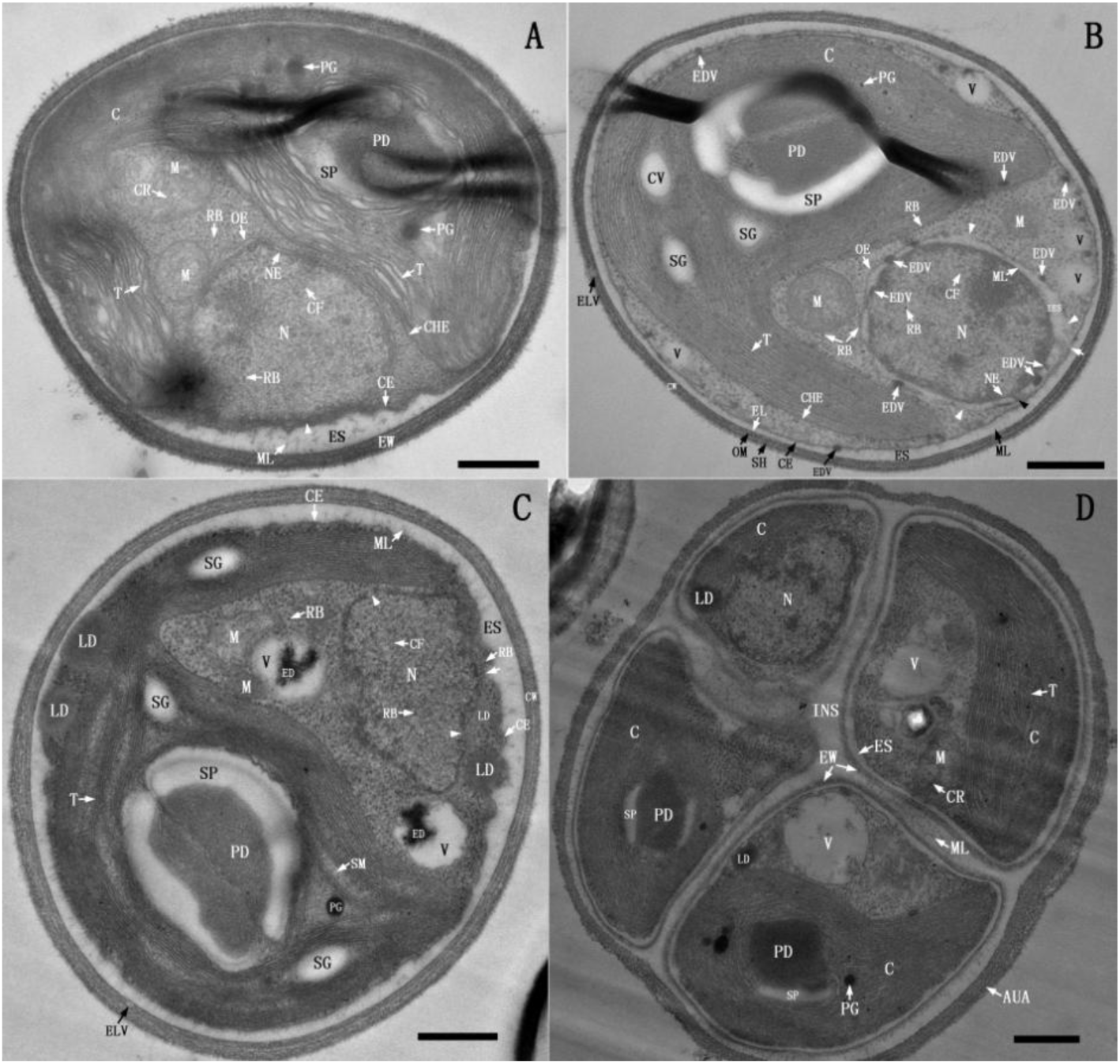
Structure and reproduction of TDX16-DE (day 9-10). (**A**), **(B) and (C).** ThreeTDX16-DE cells contain chloroplasts (C), nuclei (N) with chromatin fibers (CF) and ribosomes (RB), mitochondria (M) and vacuoles (v) (**(A)** had no vacuole). **(A)** The nuclear envelope (NE) and outer nuclear envelope (OE) contact the cytoplasmic envelope (CE) at the opening of chloroplast cavity (arrowhead). (**B**) Electron-dense vesicles (EDV) bud form NE (black arrowhead) into inter envelope space (IES), and then fuse with and re-bud from OE, ultimately reach the two sides of chloroplast envelope (CHE) and CE. There are several openings on OE (white arrowhead), and one opening at the contact site of OE and CE (arrow). (**C**) OE and NE contact intimately, on which large openings (arrowhead) are formed, and also a fusion pore is developed at the contact site of NE, OE and CE (arrow). (**D**) Four autospores in an autosporangium (AUA) are segregated from each other by the wide interspace (INS). Scale bar, 0.5 μm.

TDX16-DE multiplies via autosporulation. As shown in Fig.14D, four autospores (TDX16-DE cells) within an autosporangium (AUG) are in different developmental stages and more or less similar in arrangement to TDX16 (endospores) in the sporangium (Fig.2).

### Photosynthetic pigments of TDX16 and TDX16-DE

In vivo absorption spectra (Fig.15A) showed that apart from the absorption maxima of chlorophyll a (Chl a) at 440 and 680 nm, TDX16 displayed a prominent peak at 630nm, corresponding to phycocyanin [51]; while TDX16-DE cell lacked phycocyanin peak at 630nm but exhibited a conspicuous shoulder peak of chlorophyll b (Chl b) at 653 nm [52], and a merged peak of carotenoids around 485 nm. Consistently, fluorescence emission spectroscopy indicated that the water soluble pigment extract of TDX16 (Fig. 15E) and lipid soluble pigment extract of TDX16-DE (Fig. 15F) displayed an emission peak of phycocyanin at 646 nm [53] and an emission peak of Chl b at 658 nm [54] respectively, but no emission peak was detected in the water soluble pigment extract of TDX16-DE (Fig. 15E) and lipid soluble pigment extract of TDX16 (Fig. 15F). The separated phycocyanin of TDX16 showed an absorption peak at 617nm (Fig. 15B), nearly the same as that of C–phycocyanin [53]; the purified Chl b and lutein of TDX16-DE displayed absorption peaks at 456 and 645 nm (Fig. 15C), 420, 446 and 475nm (Fig. 14D) respectively, identical to those of plant pigments [55]. These results demonstrate that TDX16 contains phycocyanin but no Chl b, while TDX16-DE has Chl b but no phycocyanin.

**Figure 15.**
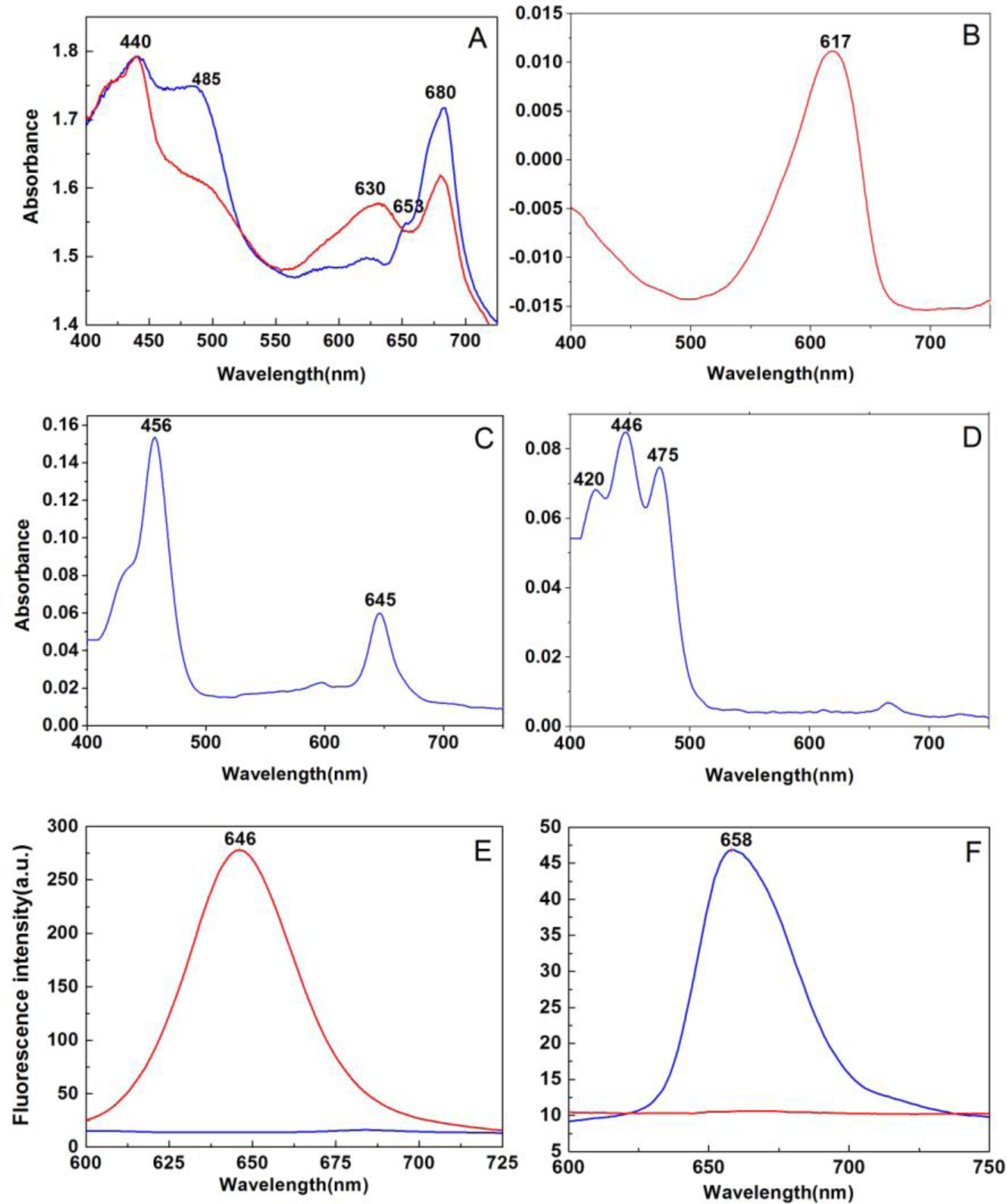
Absorption and fluorescence emission spectra. (**A**) In vivo absorption spectra of TDX16 (red) and TDX16-DE cell (blue). Absorption spectra of the isolated and purified phycocyanin (**B**), chlorophyll b (**C)** and lutein (**D**). Fluorescence emission spectra of water soluble pigment extracts (**E**) and lipid soluble pigment extracts (**F**) of TDX16 (red) and TDX16-DE cell (blue).

### 16S rRNAs of TDX16 and TDX16-DE chloroplast

16S rRNAs of TDX16 (GenBank KJ599678.2) and TDX16-DE chloroplast (GenBank KJ612008.1) share a low identity of 83%, but show high similarities of 98% and 99% to that of *Chroococcidiopsis thermalis* (GenBank NR102464.1) and those of the chloroplasts of *Auxenochlorella protothecoides* (GenBank AY553213.1) and *Chlorella vulgaris* (GenBank AB001684.1) respectively.

### TDX16 genome

TDX16 genome is 15,333,193 bp in size with an average of GC content of 55.2 % containing 15,894 genes (CDS 15,756; RNA 138). This Whole Genome Shotgun project has been deposited at DDBJ/ENA/GenBank under the accession NDGV00000000. The version described in this paper is version NDGV01000000.

## Discussion

### The reason for organelle biogenesis in TDX16

Organelle biogenesis in prokaryotes is reasonable in theory. However, the formation of organelles in TDX16 may be intuitively unacceptable and suspected as an artifact of contamination with microorganisms, because (1) such an event had not been observed previously, and (2) it is assumed that organelle biogenesis in prokaryotes occurred only once in ancient time during the origin of the first eukaryotic common ancestor (discussion is in the following section). Indeed, the possibility of contamination by other microorganisms in this study can be completely excluded. Because (1) TDX16 cells used in the experiments were prepared from the pure and axenic colony (Fig. S3); (2) light microscopic observation confirmed the absence of other microorganisms in TDX16 cultures (Fig. 1); and (3) the intermediate cell structures between TDX16 and TDX16-DE (Fig. 1-14) are unprecedented and coherent, which can not be generated by any other microorganisms. Such that, a key question arises as to why organelles formed in TDX16 ?

The consistent results of cell morphology and color (Fig. S1 and Fig. 1A,), structure (Fig. S2 and Fig.2-3), pigmentation (Fig.15) and 16S rRNA sequence indicate that TDX16 is a phycocyanin-containing cyanobacterium resembling *C. thermalis*. However, the size, gene number and GC content of TDX16 genome (GenBank NDGV00000000) are 2.4, 2.8 and 1.2 times those of *C. thermalis* (6,315,792 bp, 5593 genes and 44.4% GC, GenBank CP003597.1), respectively. In addition, comparisons with genomes of other prokaryotes and unicellular green algae (eukaryotes) sequenced so far show that (1) TDX16 genome is the largest prokaryote genome and even larger than that of green alga *Ostreococcus tauri* (12.56MB, GenBank CAID00000000.1), and (2) gene number of TDX16 genome is much larger than those of other cyanobacteria, and larger than those of unicellular green algae, including *Chlamydomonas reinhardtii* (15,143 genes, GenBank NW_001843471), the close relative of *H. pluvialis*. These results demonstrated that TDX16 had obtained a huge number of eukaryotic genes that have higher GC contents than the prokaryotic ones. Because (1) TDX16 multiplied in the senescent/necrotic *H. pluvialis* cell at the expense of host’s degraded organelles and cytoplasmic matrix (Fig.S2), and (2) Chl b and lutein, the typical pigments of green algae, are absent in TDX16 but present in TDX16-DE (Fig. 15), it is certain that TDX16 had acquired 9,017,401bp DNAs with 10301 genes from its dead green algal host *H. pluvialis*. Accordingly, organelle biogenesis in TDX16 resulted from hybridizing the acquired DNAs with its own ones and expressing the hybrid genome. Since this study focuses on the cellular mechanism of organelle biogenesis in TDX16, we are unable to elucidate the molecular mechanism here, but infer the molecular process based on the changes of cellular structure:

1. when the bloated *H. pluvialis* cells underwent senescence/necrosis (Fig. S1), the tiny dormant endosymbiotic cyanobacterium TDX16 was activated, which took up host’s DNAs and retained the acquired DNAs most likely in the unique replicable heterogenous globular body so as to keep from degradation (Fig. S2). After the small TDX16 cells were liberated from the ruptured host cell (Fig. S1), the heterogenous globular body with electron-dense DNA-like materials situated in the nucleoid and seemed to be a “micronuleus” (Fig. 2).
2. upon compartmentalization, the obtained DNAs were released gradually from the heterogenous globular body into the nucleoid (Fig. 4), most of which kept inactive because the initially formed primary thylakoids were cyanobacterial ones (Fig.5) similar to those of *Chroococcidiopsis* [13], owing to expression of TDX16’s own genes that were suppressed within the host cell and in the dim light.
3. during de-compartmentalization the total DNAs in the solubilized inner cytoplasm fragmented, intermingled (hybridized) (Fig.6), which was then partitioned into the new intracytoplasmic space (major fraction) and new inner cytoplasm (minor fraction) during re-compartmentalization (Fig.7), and subsequently gave rise to nuclear genome (Fig.8-9) and primitive chloroplast genome (Fig.8) respectively. Hereafter, the primitive chloroplast genome partitioned into chloroplast genome and mitochondrial genome during assembly of mitochondria (Fig. 9-11). The disappearance of phycocyanin (Fig.15) and TDX16 16S rRNA, but appearance of chlorophyll b, lutein (Fig.15) and new 16S rRNA of TDX16-DE chloroplast indicated that DNA hybridization resulted in the loss of prokaryotic genes, retention of eukaryotic genes and synthesis of new (hybrid) genes.

### Organelle biogenesis in TDX16 sheds light on eukaryotes

Organelle biogenesis in the prokaryote TDX16 (Fig. 2-13) resulted in its transition into an eukaryote TDX16-DE (Fig. 14), that is to say, the organelles of eukaryote TDX16-DE were originally formed in the prokaryote TDX16. So, the biogenesis of organelles in TDX16 provides a reference for re-understanding the development, structure, function and association of organelles and other compartments in eukaryotes and the reasons behind them.

#### Formation of different DNA-containing organelles all at once

With the initial compartmentalization (Fig. 4) and subsequent de- and re-compartmentalization (Fig.6-7), the whole cellular content of TDX16 was allocated into the only two compartments: the new intracytoplasmic space and the new inner cytoplasm (Fig.7), from which nucleus, chloroplast and mitochondrion were developed respectively. (Fig.8-11). Therefore, different DNA-containing organelles in TDX-DE were formed all at once in a single development event with concurrent establishment of interorganellar metabolic link and communication as evidenced by the viability of TDX16-DE. Accordingly, the interorganellar signaling (control) is a multilateral system, encompassing not only anterograde signaling (nucleus-to-chloroplast/mitochondrion) and retrograde (chloroplast/mitochondrion-to-nucleus) signaling [56–58], but also chloroplast-to-mitochondrion signaling [59] and mitochondrion-to-chloroplast signaling.

#### The number of membranes enclosing the DNA-containing organelles

Chloroplast, nucleus and mitochondrion were all formed in TDX16 by encapsulating the relevant components with the membranes synthesized by fusion, flatten and extension of the small vesicles (Fig. 6-10). Such that, the envelopes of these DNA-containing organelles inevitably comprise two unit membranes if the vesicles are bounded by one unit membrane, but four unit membranes, such as the nuclear envelop of TDX16-DE (Fig.14) and the chloroplast envelopes of some algae [60–62], if the vesicles are bounded by two unit membranes.

#### Uneven DNA distribution and promiscuous DNA sequences among organelles

During de- and re-compartmentalization the total DNAs in TDX16 aggregated (hybridized) and were partitioned into the new intracytoplasmic space (major fraction) and the new inner cytoplasm (minor fraction) (Fig.6-7). The former’s matrix was then encapsulated into the nucleus (Fig.8), while the latter subsequently gave rise to chloroplast and mitochondrion (Fig.8-12). So, nucleus sequestered most of the cellular DNAs, while chloroplast and mitochondrion contained only a handful of DNAs. Accordingly, DNA partition during organelle biogenesis in prokaryotes is the cause for uneven DNA distribution among organelles in eukaryotes, and likely also the cause for promiscuous DNA sequences: chloroplast and mitochondrial sequences in nucleus [63], nuclear and chloroplast sequences in mitochondrion [64–66], and nuclear and mitochondrial sequences in chloroplast[67–71], which are usually interpreted as interorganellar, intracellular or horizontal (DNA/gene) sequence transfers.

#### The common origin of eukaryotic cytoplasmic matrix and nuclear matrix

After formation of the two transitional composite organelles, the eukaryotic cytoplasmic matrix was built up from the matrix extruded by the primitive nucleus (Fig.9), indicating the common origin of eukaryotic cytoplasmic matrix and nuclear matrix (nucleoplasm). Therefore, it is not surprising that (1) most of nucleus-encoded proteins are synthesized in the eukaryotic cytoplasmic matrix, (2) some nucleus-encoded proteins are synthesized locally in the nuclear matrix of TDX16-DE as indicated by the presence of ribosome (Fig.14) and the nuclei of other cells (nuclear translation) [72–77], and (3) during open mitosis, (A) nucleus but not mitochondrion or chloroplast disassembles, (B) nuclear matrix and eukaryotic cytoplasmic matrix coalesce into one compartment (the reverse process of their formation), and (C) daughter nuclei form by de novo assembly of nuclear envelop (in this regard, daughter nuclei form de novo). Accordingly, the eukaryotic cytoplasmic matrix and nuclear matrix is in essence a dynamic entity, and thus we define them as the nucleocytoplasmic matrix.

#### Development of mitochondria in the primitive chloroplasts

Mitochondria were assembled in and segregated from the primitive chloroplast in TDX16 (Fig. 9-11), demonstrating that mitochondria and chloroplasts developed from the same intermediate organelle, which is the reason why (1) chloroplast and mitochondrial genomes share similar features [78], and have widespread homologies and some same DNA sequences [79–80], (2) mitochondria contain specific chloroplast proteins [81], and (3) chloroplasts possess respiratory electron transfer chains for chlororespiration [82–83].

The presence of mitochondria in chloroplasts [84–89] and detachment of mitochondria or mitochondrion-like bodies from chloroplasts [90–92] have been observed frequently in plant cells. However, these phenomena have largely been overlooked and whether the chloroplast-localized mitochondria were developed in or engulfed by the chloroplast remains controversial. In light of the assembly of mitochondria in the primitive chloroplast of TDX16, it is likely that the impaired mitochondria entered the chloroplast and dismantled, allowing the mixture of mitochondrial matrix and chloroplast stroma (the reverse process of their formation). Such that the chloroplast became a primitive-chloroplast-like composite organelle, in and from which new mitochondria were assembled and segregated. Thus, assembly of mitochondria in chloroplast is, in principle, similar to development of daughter nuclei in nucleocytoplasmic matrix during open mitosis, which occurs in the plant cells that probably contain only impaired mitochondria under stress conditions.

#### Transition of mitochondria into double-membraned vacuoles

Mitochondria in TDX16 turned into double-membraned vacuoles after matrix degradation (Fig.11). Currently, mitochondrion in plant cells is thought to be degraded only by autophagy (mitophagy), which is selectively enclosed by the phagophore (isolation membrane) into a double-membraned autophagosome, and then delivered into the single-membraned vacuole for degradation. This way of mitochondrion degradation is, however, unachievable in TDX16 cell, because (1) autophagosome can not be formed owing to the lack of endoplasmic reticulum, which is proposed to be the membrane source for phagophore synthesis in plant cells[93], and (2) there is no vacuole, which is also developed from scratch.

The cellular mechanism of vacuole biogenesis in plant cell is unclear, while the vacuolar membrane is presumably sourced from Golgi apparatus[94] or endoplasmic reticulum[10], both of which are devoid in TDX16 and TDX16-DE. So, mitochondrion-to-vacuole transition in TDX16 is an alternative way of mitochondrion degradation and vacuole formation, while the hydrolytic enzymes were probably stored in and released from mitochondria’ internal bodies (Fig.10-11). Since double-membraned vacuoles also present in plant cells [95–96], it is certain that transition of mitochondria or other organelles (e.g., autophagosomes) into double-membraned vacuoles occur in plant cells under specific conditions.

Different from the single-membraned vacuoles in plant cells, the double-membraned vacuoles in TDX16-DE serve both as a degradative and as a secretory compartment, similar in function to the secretory lysosomes in animal cells [6–7]. Since mitochondria-produced vesicles contribute to the generation of peroxisomes [5], it is possible that the mitochondria-derived double-membraned vacuoles in TDX16-DE also play the roles of peroxisomes.

#### Formation of different thylakoids

Thylakoids are photosynthetic apparatus and vital for cell maintenance and organelle development. So, concomitant with compartmentalization, de- and re-compartmentalization were the assembly/disassembly of primary and secondary thylakoids in the transitional compartments and finally development of primitive eukaryotic thylakoids in the primitive chloroplast (Fig.4-8).

1. The primary thylakoids developed from the electron-transparent vesicles formed by “germination” of osmiophilic granules (Fig.4-5). Thus, (A) osmiophilic granules are the precursors of thylakoids in cyanobacteria, which is why there is no osmiophilic granule in the thylakoid-less cyanobacterium *G. violaceus* [30]; (B) osmiophilic granules consist of two rather than one half unit membrane; and (C) the intralamellar vesicles of cyanobacteria [97–99] are premature thylakoids developed from osmiophilic granules.
2. The secondary thylakoids developed from the primary thylakoid-derived vesicles (Fig.7), in a manner similar to re-development of thylakoids from the former thylakoid-derived vesicles or segments in the stress-experienced cyanobacterial cells and germinated akinetes [17,100]. Nevertheless, the absence of phycobilisomes in secondary thylakoids suggested that the primary thylakoid-derived vesicles were modified in composition.
3. The primitive eukaryotic thylakoids developed from the plastoglobuli produced during disassembly of secondary thylakoids in the same way as the development of primary thylakoids from the osmiophilic granules (Fig.8). Therefore (A) plastoglobuli are identical to osmiophilic granules in structure but different in composition, and (B) plastoglobuli are the precursors of chloroplast thylakoids, which is why their number increases during thylakoid breakdown but decreases during thylakoid development [25, 101].

#### The double-membraned cytoplasmic envelope of TDX16-DE

TDX16-DE’s double-membraned cytoplasmic envelope was formed in TDX16 by combination of the outer intracytoplasmic membrane with the cytoplasmic membrane after degradation of the outer cytoplasm (Fig. 4), the two membranes of which were closely appressed in most cases and thus difficult to distinguish without being aware of the developmental process.

As a matter of fact, double membraned cytoplasmic envelopes seem also present in other unicellular green algae, particularly picoalgae (cell size less than 3 µm), which are ignored or thought to be the two layers of a unit membrane. For examples, *Nanochlorum eucaryotum*, *Nannochloris coccoides* [102], *Chlorella fusca* [42] and *Chloroparva pannonica* [103] apparently have double-membraned cytopalsmic envelops akin to that of TDX16-DE, while *Pseudochloris wilhelmii*[104], *Nannochloropsis oceanica*[105], and *Chlorella sp*[106–108] seem also to be the case. Picoalgae are very simple in structure, containing a minimal set of organelles. For examples, the picoalgae [109–117] all lack endoplasmic reticulum, most of which also lack Golgi apparatus [109-111, 114-115], and few of which lack endoplasmic reticulum, Golgi apparatus and peroxisome [116–117] and even vacuole [117]. These results suggest that endoplasmic reticulum, Golgi apparatus and peroxisome are not essential and their functions can be performed in other compartments, similar to performance of mitochondrial function in cytoplasmic matrix of the mitochondrion-less eukaryote *Monocercomonoides* sp[118].

TDX16-DE is very small (2.9-3.6 µm) and bears strong resemblances to picoalgae in cell morphology, structure, organelle number and arrangement, though 16S rRNA sequence of its chloroplast shows high similarity to those of the chloroplasts of *A. protothecoides* and *C. vulgaris*. In TDX16-DE cells, the attachment of ribosomes to and assembly of lipid droplets on the inner membrane of cytoplasmic envelop (Fig.10-14) indicate that the cytoplasmic envelop serves the functions of endoplasmic reticulum for synthesizing proteins and lipids. And the proteins synthesized on the inner membrane can be used locally for remodeling the cytoplasmic envelop without sorting and secreted into the extracytoplasmic space directly with no need for vesicular transport, and thus the cytoplasmic envelop serves the functions of Golgi apparatus as well.

#### The outer membrane in the eukaryotic cell wall of TDX16-DE

The eukaryotic cell wall of TDX16-DE (Fig.8-14) was developed on the base of TDX16’s prokaryotic cell wall (Fig. 4) by forming sheath over the outer membrane and modifying the peptidoglycan layer into the electron-dense layer (Fig.4D-8A), and so, the outer membrane remains in the eukaryotic cell wall of TDX16-DE. Since the sheath scaled off and re-formed continuously until formation of the stratified sheath, the outer membrane where enzymes anchored was remodeled to accommodate the above changes, which along with the electron-dense layer and the sandwiched electron-transparent space constituted a trilaminar domain resembling the trilaminar sheaths [41–45, 106] in algal and plant cell walls. The trilaminar domain and the trilaminar sheaths are equivalent structures, because like cyanobacterial cell walls, algal and plant cell walls also contain lipids [43,119-122], enzymes[43-44, 123-126] and carotenoids [127–128]. Accordingly, the out layers of trilaminar sheaths in algal and plant cell walls are membranes, which serve as platforms for anchoring enzymes to assemble the primary and secondary cell walls (corresponding to the sheath and the extended part of electron-dense layer in the eukaryotic cell wall of TDX-DE, respectively) and barriers for constraining lateral diffusion of cytoplasmic membrane proteins [129]; while the intermediate layers are aqueous spaces and the inner layers are porous mechanical structures.

#### The extracytoplasmic space is a membrane-surrounded compartment

TDX16 had an extracytoplasmic space between the cytoplasmic membrane and the outer membrane (Fig. 4A), equivalent to the periplasmic space of other prokaryotic cells [27, 130]. The extracytoplasmic space remains in TDX16-DE (Fig.14), which is surrounded by the cytoplasmic envelope and the outer membrane within the eukaryotic cell wall, and thus a genuine membrane-surrounded compartment. During biogenesis of the primitive nucleus the liquid of new intracytoplasmic matrix was squeezed out into the extracytoplasmic space (Fig.8), and later the vacuoles expelled their contents into and internalized substances from it (Fig.13). So, the extracytoplasmic space plays indispensable roles and metabolically links to the eukaryotic cell wall, cytoplasmic envelope, eukaryotic cytoplasmic matrix, nucleus and vacuoles.

Likewise, the extracytoplasmic spaces (interspaces) between the cytoplasmic membrane and the trilaminar-sheath-containing cell wall in algal cells[41–42, 44] are also membrane-surrounded compartments, rather than the extracellular spaces (apoplasts) [131] as usually considered.

#### Generation of different small bodies from the primitive chloroplast

The oversized primitive chloroplast in TDX16 was the source of energy and materials for organelle development and cell maintenance, from which different small bodies generated during it dwindled into a large chloroplast.

1. the vesicle-containing body was firstly developed from the primitive chloroplast and engulfed by the primitive nucleus (Fig.9A), which delivered materials for primitive nucleus maturation and formation of eukaryotic cytoplasmic matrix. The absence of visible enclosing-membrane and rectangular-shape of the vesicle-containing body suggested that it was formed most likely by incision, but not by budding or protrusion.
2. the chloroplast debris with fragmented primitive thylakoids (Fig. 11B, 12E and 13D) was irregular in shape and devoid of enclosing-membrane, and thus seemed to be chipped off the primitive chloroplast during assembly of mitochondria and degraded, at least in part, in the eukaryotic cytoplasmic matrix, which originated from the primitive nucleus that was capable of degrading the vesicle-containing body.
3. the compound vesicles detached from the primitive chloroplast in a way similar to that of mitochondria and transferred principally the segmented primitive eukaryotic thylakoids into vacuoles for degradation and formation of multilamellar bodies (Fig. 12).

These small bodies played roles in primitive chloroplast maturation, the mechanisms by which they formed apparently differed from those for developing the small bodies during the piecemeal-degradation of chloroplast, including Rubisco-containing body [132], ATI1-PS body [133] and SSGL body [134], as well as senescence-associated vacuole [135] and CV-containing vesicle [136]. Hence, generating different small bodies through different mechanisms is a strategy for both chloroplast development and chloroplast degradation.

### Implications of organelle biogenesis in TDX16

This study unveils the biogenesis of organelles in cyanobacterium TDX16. Likewise, it is reasonable that organelles can also form in bacteria in the similar situations. In light of organelle biogenesis in TDX16, we postulate that organelles (no chloroplast) form in the eukaryotic-DNA-acquired bacteria by enclosing the cytoplasm into a primitive nucleus, from which mitochondria and eukaryotic cytoplasmic matrix develop. This postulation is supported by the facts that in heterotrophic eukaryotes (1) mitochondria present in nuclei [137–142], (2) nuclear genomes contain mitochondrial DNA copies [143–145], and (3) most mitochondrial proteins are descendants of nuclear genes with no bacterial antecedents [146–148]. Therefore, organelle biogenesis in TDX16 has broad implications on biology, particularly cancer biology and evolutionary biology.

#### Implications on cancer biology

The origin of cancer cells is the most fundamental yet unresolved problem in cancer biology. Cancer cells are thought to be transformed from the normal cells, however, recent studies reveal that the small nascent primary cancer cells (PCCs) for cancer initiation [149–153] and secondary cancer cells (SCCs) for cancer progression [151, 154-162] are formed in and but not transformed from the senescent normal cells and cancer cells, respectively. These small nascent PCCs/SCCs are very small, undifferentiated [154, 161, 163-164] and absent of organelles, which mature into eukaryotic PCCs/FCCs after being released from the ruptured senescent normal/cancer cells. Nevertheless, the cellular mechanisms of how the small nascent PCCs/SCCs formed, and how organelles developed in PCCs/SCCs are unclear. In the light of TDX16-to-TDX16-DE transition, it is most likely that PCCs/SCCs arise from bacteria [165]: the intracellular bacteria take up the senescent normal/cancer cells’ DNAs and become small nascent organelle-less PCCs/SCCs, which develop into eukaryotic PCCs/SCCs by hybridizing the acquired DNAs with their own ones and expressing the hybrid genomes to guide organelle biogenesis.

#### Implications on evolutionary biology

The origin and diversification of organelles are two different problems in evolutionary biology. The origin of organelles is coupled to origin of eukaryotes concerning how the ancestral organelles formed in the first and last eukaryotic common ancestors. This problem is difficult to make clear owing to the lack of direct evidence. Now, the endosymbiotic hypothesis [166–167] is widely accepted that an endosymbiotic eubacterium and an endosymbiotic cyanobacterium within an eukaryotic cell turned into the ancestral mitochondrion and chloroplast respectively, while an endosymbiotic archaebacterium within an eubacterium transformed into the ancestral nucleus [168–170]. By contrast, the diversification of organelles is coupled to formation of new single-celled eukaryotes (the progenitors of multicellular eukaryotes) regarding how the new organelles form in the new single-celled eukaryotes after the formation of the first and last eukaryotic common ancestors. This problem has received little attention and remains unclear.

Organelle biogenesis in TDX16 (1) uncovers a way of organelle diversification and new single-celled eukaryote formation: the endosymbiotic prokaryote acquires and hybridizes its eukaryotic host’s DNA with its own one and then develops into a new single-celled eukaryote by de novo biogenesis of new organelles; (2) provides clues for inferring the origin of organelles and modifying the endosymbiotic hypothesis. In light of organelle biogenesis in TDX16, it is more likely that the ancestral nucleus, mitochondrion and chloroplast were de novo formed in but not transformed from the endosymbiotic prokaryotes that had acquired their senescent/necrotic hosts’ DNAs. Such that, the modified endosymbiotic theory unifies the endosymbiotic theory and autogenous theory, and accounts for both the origin and the diversification of organelles.

## Methods

### Strain and cultivation

Pure TDX16 cells were collected from the *H. pluvialis* cultures, in which all *H. pluvialis* cells burst and released TDX16 cells[11], and maintained in BG-11 liquid medium [171] at 25°C, 12 μmol photons m^−2^ s^−1^ in the illumination incubator.

Axenic TDX16 was prepared according to the method [172] with some modifications. Briefly, pure TDX16 culture was treated with antibiotics nystatin (100 μg/ml) and cycloheximide (150 μg/ml) for 18h, then the culture was diluted 10 fold with sterile distilled water and plated onto BG-11 solid medium in petri dishes supplemented (w/v)with glucose (0.5 %), peptone (0.3 %) and yeast extracts (0.2 %). The petri dishes were sealed with Parafilm and incubated in an inverted position in the illumination incubator at 25 °C, 12 μmol photons m^−2^ s^−1^ for one month. The colonies (Fig. S3) were picked and transferred into autoclaved 250-ml flasks containing 100 ml BG-11 medium and cultivated under the same conditions as described above.

The obtained axenic TDX16 cultures were used in this experiment. For TDX16 transition, 10 ml axenic TDX16 culture was inoculated into each autoclaved 250-ml flask containing 100 ml BG-11 medium and illuminated with continuous light of 60 μmol photons m^−2^ s^−1^ at 25 °C.

### Microscopy observations

#### Light microscopy

TDX16 cells in cultures were examined each day with a light microscope BK6000 (Optec, China). Photomicrographs were taken under the oil immersion objective lens (100×) using a DV E3 630 camera. Cell sizes were measured with a micrometer eyepiece.

#### Transmission electron microscopy

TDX16 cells were harvested each day by centrifugation (3000 rpm, 10 min) and fixed overnight in 2.5% glutaraldehyde (50 mM phosphate buffer, pH7.2) and 1% osmium tetroxide (the same buffer) for 2 h at room temperature. After dehydration with ascending concentrations of ethanol, the fixed cells were embedded in Spurr’s resin at 60°C for 24 h. Ultrathin sections (60 to 80 nm) were cut with a diamond knife, mounted on a copper grid, double-stained with 3% uranyl acetate and 0.1% lead citrate, and examined using a JEM1010 electron microscope (JEOL, Japan).

### Pigment analyses

#### In vivo absorption spectra

Cell suspensions were scanned with Ultraviolet-Visible Spectrophotometer Cary 300 (Agilent, USA), the spectra were normalized to give an equal absorbance of Chl a at 440 nm.

#### Fluorescence emission spectra

Water soluble pigments were extracted with 0.75M K-phosphate buffer (pH=6.8). Lipid soluble pigments were extracted with pure acetone and diluted 50-fold into ethanol. Both extracts were analyzed on Fluorescence Spectrophotometer F-4500 (Hitachi, Japan) at room temperature with excitations of 580 nm and 478 nm respectively.

#### Pigment separation and identification

Chl b and lutein was separated by thin-layer chromatography according to the method described by Lichtenthaler [55]. phycocyanin was extracted and purified following the procedures described by Adams [173]. All pigments were analyzed with Ultraviolet-Visible Spectrophotometer Cary 300 (Agilent, USA), and identified by spectroscopic matching with the published data.

### 16S rRNA sequence

DNA samples were prepared according to the method described previously [174]. 16S rRNAs were amplified using the primers 8-27f (AGAGTTTGATCCTGGCTCAG) and 1504-1486r (CTTGTTACGACTTCACCCC) [175]. Fragments were cloned into the pMD18-T vector and amplified using M13 forward and reverse universal primers. The PCR products were digested with restriction enzymes BamH1/SalI, and sequenced on ABI 3730 DNA analyzer (PerkinElmer Biosystems, USA).

### Genome sequence of TDX16

TDX16 cells were harvested by centrifugation at 3000 rpm for 10 min, and washed twice with 5M NaCl solution and sterile water alternately. The pelleted cells were frozen in liquid nitrogen and then grinded with sterile glass beads (0.5 mm diameter). The slurry was transferred into 5ml centrifuge tube with TE buffer (1mM EDTA, 10 mM Tris-HCl, pH=8.0), supplemented with 1.0 ml lysozyme solution (20 mg/ml) and incubated at 37 °C for 60 min, then added CTAB (Cetyltrimethyl ammonium bromide) solution (10% CTAB, 0.7 M NaCl) and heated to 65°C for 30 min in a water bath. After centrifugation (12000 rpm, 10min), the supernatant was extracted with one volume of phenol-chloroform-isoamyl alcohol (25:24:1, V/V), and DNA was precipitated overnight at −20 °C after the addition of 2/3 volume of cold isopropanol and 1/10 volume of 3M sodium acetate, dried and resuspended in TE buffer. The extracted DNA was first subjected to quality assay and then sheared ultrasonically into fragments, with their overhangs being converted into blunt ends applying T4 DNA polymerase, Klenow Fragment and T4 Polynucleotide Kinase. Subsequently, adapters were ligated to the 3’ ends of DNA fragments that were introduced with ‘A’ bases. The obtained fragments were purified via gel-electrophoresis and PCR-amplified for preparing the sequencing library of DNA clusters. Paired-end sequencing was carried out on an IIlumina HiSeq 4000 platform, yeilding1.132 Mb raw data. After removal of the low quality reads, 1.009 Mb clean data was assembled with SOAPdenovo.

## Acknowledgements

This work was supported by the Natural Science Foundation of Hebei Province (B2008000029).

## List of abbreviations

AUG: Autosporangium
C: Chloroplast
CD: Chloroplast debris
CE: Cytoplasmic envelope
CF: Chromatin fibers
CG: Cyanophycin granules
CHE: Chloroplast envelope
CLM: Cloudlike materials
CM: Cytoplasmic membrane
CPV: Compound vesicles
CR: Cristae
CV: Combined vesicles
CW: Cell wall
CX: Carboxysomes
DF: DNA Fibers
DGV: Dense-margined vesicles
DLF: DNA-like fibrils
DMF: Double-layered membrane fragment
DMS: Double-layered membrane segment
DMV: Double-membraned vesicles
DRV: Dilated ring-shaped vesicles
DSV: Dense vesicle
DT: DNA threads
DV: Dotted vesicles
ED: Electron-dense debris
EDV: Electron-dense vesicles
EF: Electron-dense fibrils
EG: Electron-dense granules
EIS: Empty Inner space
EL: Electron-dense layer
ELM: Electron-translucent materials
ELV: Electron-translucent vesicles
EM: Eukaryotic cytoplasmic matrix
EOB: Electron-opaque bodies
EOP: Electron-opaque particles
EOM: Electron-opaque materials
EOV: Electron-opaque vesicles
EP: Electron-dense particles
EPM: Electron-transparent materials
ER: Endoplasmic reticulum
ES: Extracytoplasmic space
EV: Electron-transparent vesicle
EW: Eukaryotic cell wall
FM: Fibrillar materials
GA: Golgi apparatus
GP: Globular particles
HGB: Heterogenous globular bodies
IB: Intranuclear body
ICE: Intracytoplasmic envelope
ICP: Inner cytoplasm
IES: Inter envelope space
IIM: Inner intracytoplasmic membrane
IIS: Inner intracytoplasmic space
INS: Interspace
IS: Intracytoplasmic space
ITB: Internal body
IV: Internal vesicle
IVS: Invaginated space
LD: Lipid droplet
LDB: Less electron-dense bodies
LDM: Less electron-dense materials
LM: Limiting membrane
M: Mitochondrion
ME: Mitochondrial envelope
MF: Membrane fragments
ML: Microfibrils
MLB: Multilamellar body
MR: Margin residues
MS: Membrane segments
MT: Membranous elements
MV: Microvesicles
N: Nucleus
NE: Nuclear envelope
NIC: New inner cytoplasm
NIS: New inner intracytoplasmic space
NS: New intracytoplasmic space
NT: Nucleoid-like structure
NU: Nucleoid
NX: New intracytoplasmic matrix
OCP: Outer cytoplasm
OE: Outer nuclear envelope
OG: Osmiophilic granules
OIM: Outer intracytoplasmic membrane
OIS: Outer intracytoplasmic space
OPV: Opaque-periphery vesicle
OM: Outer membrane
OV: Oblong vesicles
P: Peptidoglycan layer
PB: Polyphosphate bodies
PC: Primitive chloroplast
PCB: Phycobilisomes
PD: Pyrenoids
PG: Plastoglobuli
PL: Peptidoglycan-like layer
PMT: Primitive thylakoids
PN: Primitive nucleus
PNE: Primitive nuclear envelope
PO: Pores
PT: Primary thylakoids
RB: Ribosomes
RM: Residual membranes
RV: Ring-shaped vesicles
SA: Sporangium
SG: Starch granules
SH: Sheath
SMV: Smaller vesicles
SM: Stroma
SOV: Small opaque vesicle
SP: Starch plate
ST: Secondary thylakoids
SV: Small vesicles
T: Thylakoids
TL: Thylakoid-like structure
TMF: Two-layered membrane fragment
TMV: Thick margin vesicle
TV: Tiny vesicles
V: Vacuole
VB: Vesicle-containing body

---

**Fig. S1.**
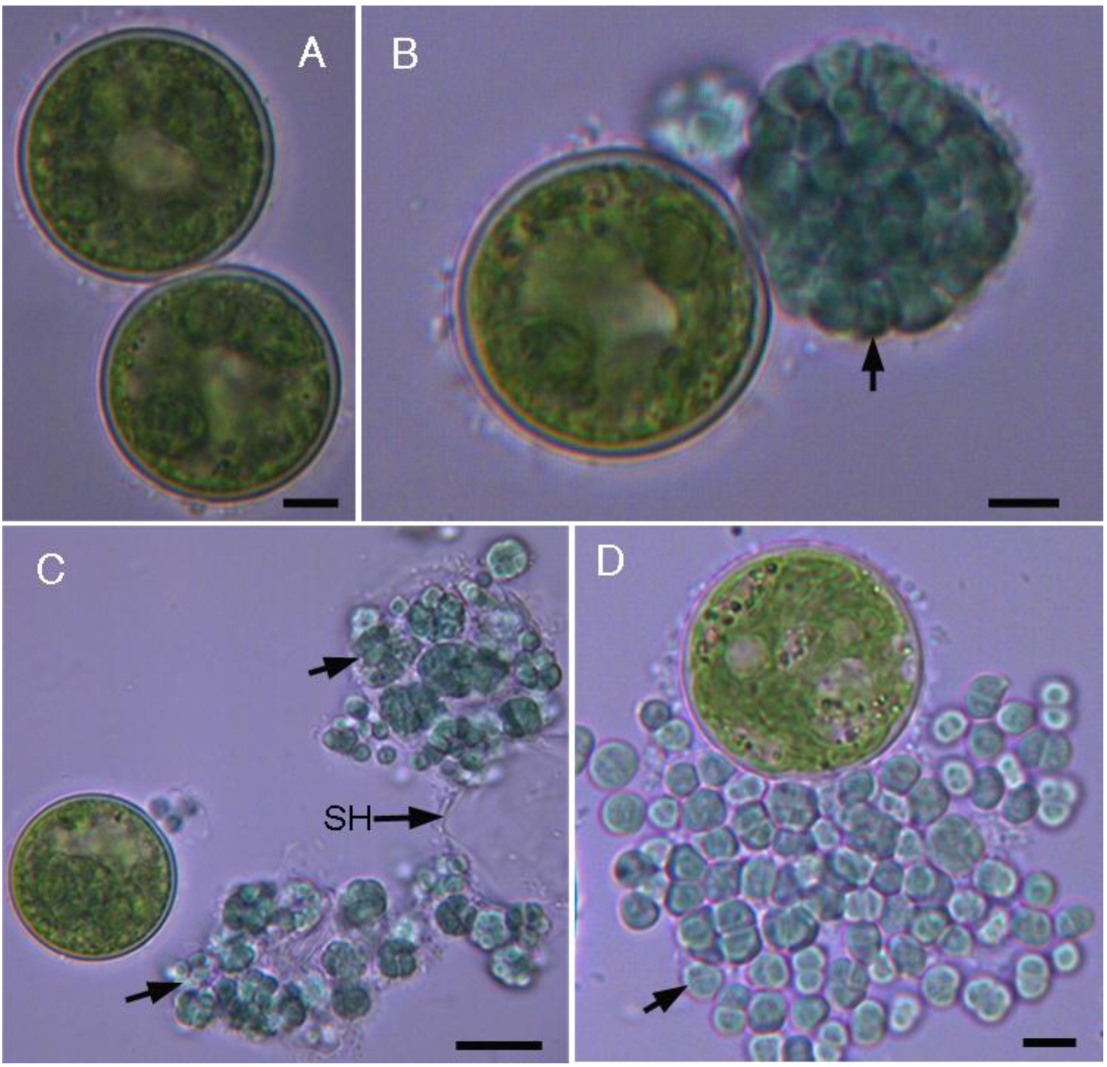
Light microscopic images of TDX16 liberation from the senescent *Haematococcus pluvialis* cell. **(A)** Two large senescent *H. pluvialis cells*, scale bar 5μm. **(B)** One senescent *H. pluvialis* cell suddenly burst and liberated a massive equal-sized blue spheroid (arrow) consisting of countless TDX16 cells, scale bar 5μm. **(C)** The transparent covering sheath (SH) of the blue spheroid ruptured, and thus the blue spheroid collapsed into many cell clumps (arrow), scale bar 10μm. **(D**) The compacted TDX16 cells in the cell clumps disaggregated, which were enclosed within the sporangia (arrow), scale bar 5μm. Cells were observed with a light microscope BK6000 (Optec, China). Microphotographs were taken under the objective lens (40×) using a DV E3 630 camera.

**Fig. S2.**
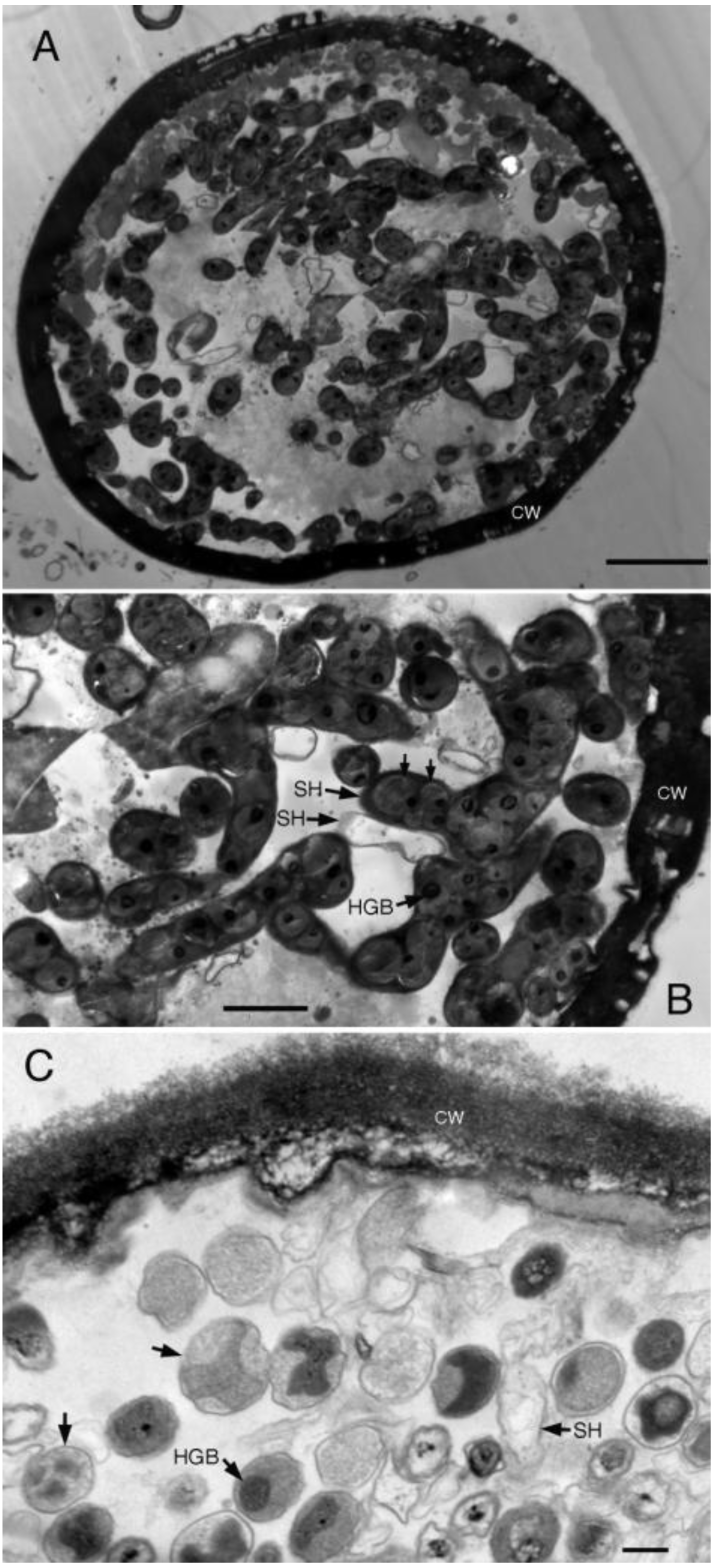
Transmission electron microscopic images of TDX16 proliferation and development in the senescent/necrotic *H. pluvialis* cell. (**A**) Tiny premature TDX16 cells proliferated within a senescent/necrotic *H. pluvialis* cell, whose organelles had dissolved, remaining only an intact cell wall (CW), scale bar 5μm. **(B)** Detail from **(A)**, tiny TDX16 cells (arrow) with electron-dense heterogenous globular bodies (HGBs) multiplied by asymmetric division within and escaped from the enclosing sheaths (sporangia) (arrow), scale bar 2μm. **(C)** Tiny TDX16 cells grew up into small thylakoid-less DTX16 cells filling up the cellular space of the dead *H. pluvialis* cell. The small TDX16 cells multiplied by formation of endospores within the sporangia (arrow), scale bar 0.2μm.

**Fig. S3.**
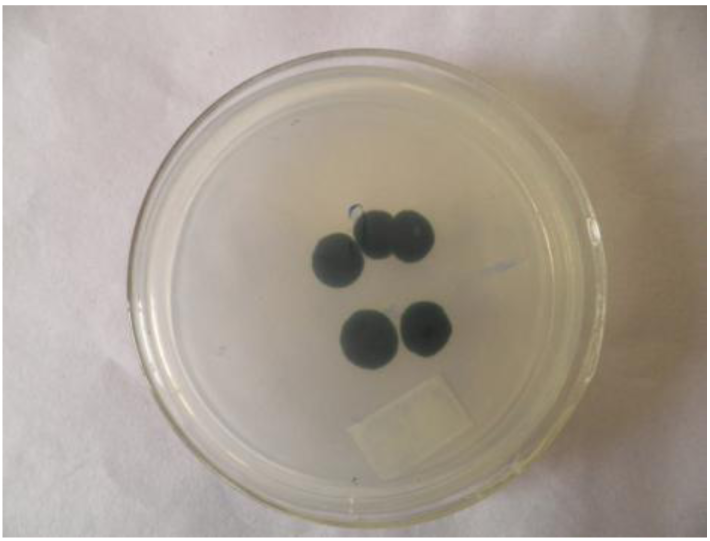
Image of TDX16 colonies in an inverted petri dish.

